# Conservation and lability within the structure of mandibular integration in “old endemic” Australian rodents

**DOI:** 10.1101/2025.06.17.660249

**Authors:** Reuben Y. Ng

**Affiliations:** Department of the Geophysical Sciences, University of Chicago, Chicago, Illinois 60637, U.S.A.

**Keywords:** phenotypic integration, modularity, rodents, Australia, cave deposits, macroevolution, developmental bias

## Abstract

The structure of phenotypic integration is predicted to bias the direction and rate of phenotypic evolution but only on timescales over which it is conserved. Both the scales on which the structure of integration evolves and how that structure is evolutionarily modified are important, but not generally understood. Here, structures of mandibular integration are inferred for eight species of “old endemic” Australian rodents, including a pair of intraspecific samples for two of these species. The structures of integration are compared and variation among these structures is assessed in light of the underlying phylogenetic relationships and in the finer patterns of conserved and evolving trait associations. Microevolution in the structure of integration is minor and while macroevolutionary comparisons almost all show significant similarity, comparisons range widely and do not clearly decay with the degree of phylogenetic separation. These patterns appear to reflect the combined influence of conserved and labile portions of the structure of integration. Structures of phenotypic integration are complexes of conserved and labile elements and if they bias phenotypic evolution, it will be in similarly complex ways.

## Introduction

While development is now well understood to bias the production of phenotypic variation available to natural selection, how (indeed, whether) this developmental bias may affect the course of evolution remains an area of active study. Covariation among traits may arise from phenotypic integration, the patterns of association between traits that arise from their developmental and functional interactions and interdependence (e.g. Cheverud, 1982, 1996; Wagner, 1996; Wagner *et al*., 2007). Phenotypic integration is predicted to promote or constrain the direction and rate of phenotypic evolution depending on the structure of integration between traits and the direction of selection (Felice *et al*., 2018). Modularity, the patterns of quasi-independence between sets of integrated traits (modules), implies a similar set of predictions (Wagner and Altenberg, 1996). Although the theory underlying phenotypic integration and the predictions that follow are increasingly well defined (see Zelditch and Goswami, 2021), these predictions have not been extensively tested and general expectations for the macroevolutionary significance of phenotypic integration (e.g. its implications regarding evolvability [Jablonski, 2022]) are in their infancy (Esteve-Altava, 2017).

Of particular importance is the stability of the structure of integration on macroevolutionary timescales because it is a predictor of patterns of disparification only on timescales over which it is conserved. The temporal/phylogenetic scale at which similarity between structures of integration decays remains essentially unknown, but almost certainly will depend on the study group, the biological structure(s) under study, the topography of the underlying adaptive landscape, and potentially the condition of the ancestral developmental system (e.g. Wagner and Altenberg, 1996; Armbruster *et al*., 2014). How the structure of integration evolves is equally important. Certain conserved elements of a structure of integration may continue to impose a bias on the production of variation, while other more labile trait associations evolve. It is therefore important not only to understand the timescales on which the structure of integration evolves, but also the patterns by which conservation begins to be lost.

The mammalian mandible has become a choice target for the study of phenotypic integration (Klingenberg and Navarro, 2012) and a great deal of work has explored how its structure of integration and modularity may relate to the many mesenchymal condensations which form the mandible (Atchley and Hall, 1991; Hall and Miyake, 2000; Hall, 2003), the pleiotropic linkages between development of different regions of the structure (Cheverud, 1996; Klingenberg and Leamy, 2001), and hypotheses of functional integration (Renaud *et al*., 2012). Here I use the mandible in eight species of “old endemic” Australian rodents as the basis for comparison between phenotypic structures of integration across micro- and macroevolutionary timescales. The primary goals of this work are (1) to quantify the timescales on which the structure of integration might serve as a persistent developmental bias, and (2) to characterise evolutionary patterns of conservation and lability within the structure of integration.

## Methods

### Data

The subfossil specimens studied here are from a remarkable series of collections of Australian cave deposits (details in supplement) housed at the Field Museum of Natural History, Chicago, U.S.A. Of the eight species studied (Figure 1A), two (*Pseudomys albocinereus* and *P. occidentalis*) are represented by both a surface sample and a subsurface sample dated to at least 7,850 *±* 170 years before present (Lundelius, 1960). Locality and sample details can be found in the supplement.

**Figure 1:**
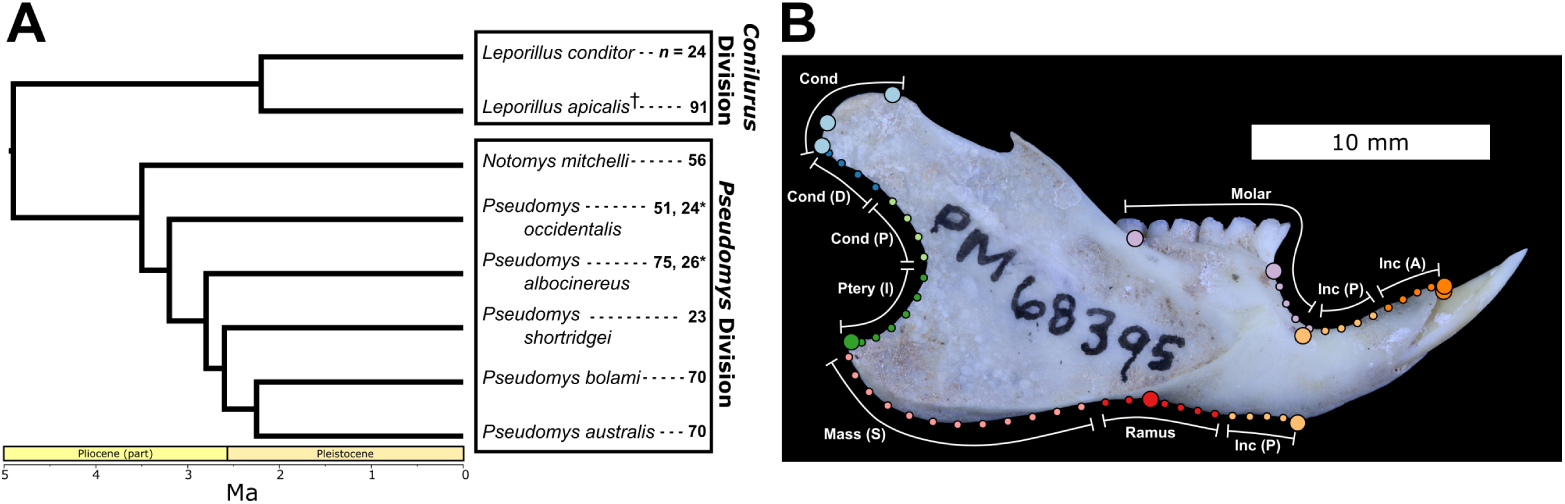
**A.** Time-scaled phylogeny of taxa studied, after Roycroft *et al*. (2021); sample sizes reported at right, starred are subsurface samples while all others are surface collections. **B.** Landmark scheme on hemimandible of *Leporillus conditor* showing landmarks (large points), semi-landmarks (small points), and partitions: Cond = condyle, Cond (D) = distal condyloid, Cond (P) = proximal condyloid, Ptery (I) = insertion of the internal pterygoideous, Mass (S) = insertion of the superficial masseter, Inc (P) = posterior incisor alveolus, Inc (A) = anterior incisor alveolus.

### Shape analysis

Hemimandibles were selected for analysis based on the quality of preservation (sample size in Figure 1A). Hemimandibles were photographed in lateral (buccal) view under a standard orientation. The hemimandible was digitised using 11 landmarks and 47 semi-landmarks among 4 semi-landmark curves (Figure 1B, landmark selection and descriptions in supplement). Semi-landmark curves were initially digitised with approximately twice as many points to aid in modelling of curve shape during superimposition. Landmark configurations for each sample were placed into Procrustes superimposition (Rohlf and Slice, 1990) and semi-landmarks were slid to minimise bending energy. Semi-landmarks were then culled to their final number. Bending energy was chosen as the preferred sliding criterion over minimising Procrustes distance because, with the data in this study, the latter method yielded results unfaithful to the original curve shapes. Bending energy has recently come under scrutiny as a criterion for sliding semi-landmarks in certain analyses of integration and modularity (Cardini, 2019; Zelditch and Swiderski, 2023), but these concerns do not extend to the methods used here. Superimposition and sliding of semi-landmarks was done in R (R Core Team, 2024, version 4.4.2) using the geomorph package (version 4.0.10). Measurement error was quantified and found to be negligible (supplement).

Allometry, size-related shape variation, can be a confounding factor when assessing static (non-ontogenetic) developmental signals, such as those of modularity and integration. Except in one sample, shape variation was found to be poorly explained by size (see supplement). The sample of *Leporillus apicalis* was found to contain an appreciable degree of allometry and shape was standardised to mean size (see supplement). Size-standardisation was not performed on the other samples.

### Inferring the structure of integration

Subsetting landmark data into a number of multivariate traits (partitions) allows for the estimation of boundaries between developmental modules and the strength of integration within and between them. Various partitioning schemes and interpretations of their biological underpinnings have been evaluated for integration and modularity in the rodent mandible (e.g. Klingenberg *et al*., 2003; Zelditch *et al*., 2008; Márquez, 2008). The purpose of the present study is not to compare the efficacy of these many partitioning schemes, but to characterise the pattern of evolution of the structure of integration and the temporal and phylogenetic scales over which this has happened. To this end, the “Angular” partitioning scheme of Zelditch *et al*. (2008, 2009) has been modified for the slightly reduced set of landmark information available for study here (Figure 1B). Nine partitions are used here and have been named according to the convention of Zelditch *et al*. (2008).

The structure of integration can be inferred from the correlations between partitions (Monteiro *et al*., 2005). First, landmark configurations for each partition are independently placed into Procrustes superimposition (although see Klingenberg, 2009, p. 408 for interpretation of different superimposition strategies). Then, for each partition, a pairwise distance matrix is calculated between the shapes of that partition in all individuals. Distances are Procrustes distances, the square root of the sum of squared distances between the landmarks of two Procrustes superimposed landmark configurations (Zelditch *et al*., 2012a). Finally, pairwise correlations are calculated between the distance matrices of all partitions. The resulting matrix of pairwise correlations between partitions is the structure of integration. An extension of this method utilises a Mantel test (Mantel, 1967) to augment correlation calculations with an associated *p*-value for each pairwise matrix correlation (e.g. Zelditch *et al*., 2008, 2009; Webster and Zelditch, 2011a,b; Anderson *et al*., 2016).

Distance matrices, correlations, and associated *p*-values have routinely been calculated in the software CORIANDIS (Márquez and Knowles, 2007), however it appears that the matrix correlations returned by CORIANDIS (as of version 1.12) are calculated as correlations between all elements of the symmetric distance matrices (including the diagonals) rather than between a corresponding upper or lower triangle from each matrix. The discrepancy here is very minor, but results in slightly inflated matrix correlations in the case of the CORIANDIS method. Because of this, I opt here to calculate distance matrices in R (script in supplement) and compare them using the Mantel test as it is implemented in the R package vegan (version 2.6.10), although similar inferences follow from using CORIANDIS (version 1.12).

### Comparing structures of integration

#### Correlations between structures of integration

Two correlation structures can be compared through a Mantel test where the null hypothesis is that the two matrices being compared are no more similar than would be expected by chance. Pairwise Mantel tests are run between all correlation structures resulting in a matrix of correlations between the inferred structures of integration of each sample. The null hypothesis of non-similarity can be inverted to test whether the two matrices are no more different than would be expected by chance. The bootstrapping approach used here follows that of Webster and Zelditch (2011b, also see similar methods used by Hallgrímsson *et al*. [2006] and Hunt [2007]) and conveniently facilitates calculation of matrix repeatability for the observed correlation matrices. These tests were run in R using script written by Annat Haber (Zelditch *et al*., 2012b, chapter 12, pp. 6–7).

#### Relative eigenanalysis

Relative eigenanalysis (or relative principal component analysis) is an alternative strategy to the partition-based methods described thus far. Instead of using pre-defined partitions and examining the associations between them, relative eigenanalysis compares patterns of variance and covariance across relative eigenvalues and eigenvectors (Flury, 1985; Mitteroecker and Bookstein, 2009; Bookstein and Mitteroecker, 2014; Le Maître and Mitteroecker, 2019). The procedure gives rise to a multivariate metric for the comparison of variance-covariance matrices which is equivalent to the Riemannian distance between matrices. Riemannian distances are measured in the space of square semi-positive definite matrices and thus are non-Euclidean, but they can be indirectly plotted by principal coordinates analysis (PCoA) which provides a least-squares Euclidean approximation of their relationships. Thus relative eigenanalysis provides the tools to produce a space in which patterns of phenotypic variance and covariance can be explored. This method involves an initial principal components analysis and the selection of the leading principal components from which to perform the relative eigenanalysis at the heart of the procedure (Le Maître and Mitteroecker, 2019, p. 1384). In this case, all principal components summarising greater than 5% variance were used which amounted to the first four axes and a total of 81.3% variance summarised. Relative eigenanalysis was run using the R package vcvComp (version 1.0.2).

#### Phylogenetic structure and the evolution of phenotypic integration

Evolutionary relationships between samples (Figure 1A) can be reduced to a matrix of pairwise phylogenetic distances (measured tip to tip in millions of years). These pairwise distances can be compared to both the matrix of pairwise correlations between structures of integration (after conversion to a distance matrix, here referred to as the integration disparity matrix) and also to the matrix of pairwise (PCoA) distances between samples derived from relative eigenanalysis. These comparisons are made in several ways. First, clustering of the integration disparity matrix and PCoA distance matrix are compared to the topology of the phylogeny. Second, pairwise correlations between structures of integration, and pairwise relative eigenanalysis distances, are regressed against their corresponding phylogenetic distances allowing examination of the degree of modification to a structure of integration in light of the independent evolution that led to the structures in question. Finally, a Mantel test is used to compare the matrix of phylogenetic distances to either the integration disparity matrix or the distance matrix resulting from relative eigenanalysis. To compare the partition-based and relative eigenanalysis approaches to one another, their resulting distance matrices can be compared using these same methods. The rank order of pairwise phylogenetic distances is unaffected by uncertainty in the node age estimates (Roycroft *et al*., 2021, fig. 1) except in the case of the node subtending the two species of *Leporillus*: it is unclear how the timing of this node compares to that of the nodes within *Pseudomys*, but the degree of independent evolution between these species is on the order of other congeneric comparisons within *Pseudomys*.

#### Phylogenetic signal in trait associations

To further dissect patterns of evolution within the structure of integration, the correlation between partitions can themselves be treated as traits. These correlations are not necessarily independent of one another and so the partial correlations between partitions are first calculated. The variation among samples for a given partial correlation between partitions can then be explored in terms of phylogenetic signal. Two statistics of phylogenetic signal are calculated here, Pagel’s *λ* and Blomberg *et al*.’s *K* (Pagel, 1999; Blomberg *et al*., 2003). Both Pagel’s *λ* and Blomberg *et al*.’s *K* range from 0 to 1 with high values indicating a trait (in this case a partial correlation between partitions) varies across phylogeny in a way that is well explained by evolution by Brownian motion (Blomberg *et al*.’s *K* may also be greater than 1 indicating stronger than expected phylogenetic signal). These metrics are poorly estimated on small phylogenies (Boettiger *et al*., 2012; Münkemüller *et al*., 2012) where they struggle with Type II error (failure to reject the null of no signal). If, even on the small phylogenetic sample studied here, signal emerges, it likely reflects a genuine phylogenetic pattern in the data. In such small phylogenetic samples, relative strength of signal is better assessed by Blomberg *et al*.’s *K* which is less prone to over-inflation of values (Boettiger *et al*., 2012). Both metrics were calculated using the R package phytools (version 2.4.4).

## Results

### Comparing correlation structures

There is considerable variation between the correlation structures of the different samples. Some notes on general patterns of association between partitions follow, but see the supplement for a complete report of the correlation structure of each sample.

Only a few associations between partitions appear to be broadly conserved (Figure 2A). One of these reflects a module consisting of the ramus and posterior incisor alveolus and is very strongly detected in every sample. This combination of traits describes the curvature of the ventral margin of the mandible along the ramus and the margin of the diastema leading into the incisor alveolus. The condyle and distal condyloid are often found in association with one another, but only infrequently recovered with the proximal condyloid. A module consisting of the internal pterygoideus insertion and insertion of the superficial masseter is found in several cases. An anterior incisor alveolus and molar module is equivocally recovered in a few samples, while in others, little association is found either between these partitions or with any other. The strongest associations are generally found between adjacent partitions.

**Figure 2:**
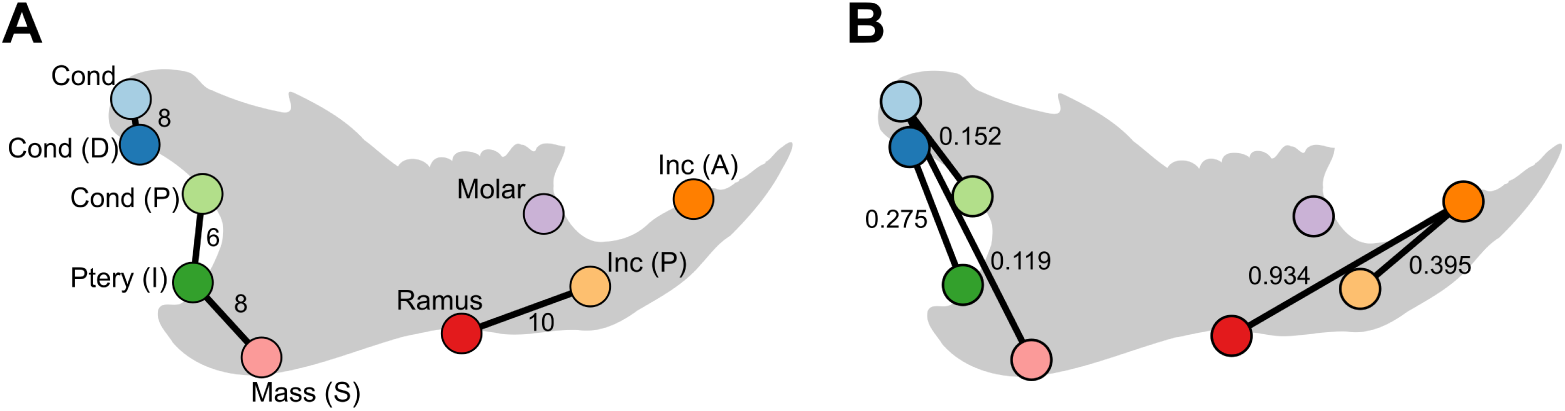
**A.** Conserved associations between partitions in six or more of the 10 samples studied (number of samples next to association); see Figure 1 for abbreviations of partitions; partitions are indicated at the mean position of their associated landmarks. **B.** Associations in which partial correlations between partitions exhibit significant phylogenetic signal; Blomberg *et al*.’s *K* reported by associations; partitions as labelled in panel **A**.

### Correlations between structures of integration

Matrix correlations between structures of integration range widely in strength (*ρ* = 0.118 *−* 0.712, Figure 3A). Most Mantel tests indicate significant similarity between structures of integration except in some comparisons involving the subsurface sample of *Pseudomys occidentalis*, *P. shortridgei*, and *Leporillus conditor*. Inverting the null of the Mantel test gave roughly a complimentary result (supplement). Cases where there is disagreement between these two tests appear to reflect correlation matrices with low repeatability and may correspond to low sample size. Conspecific comparisons lead to comparatively strong associations (*ρ* = 0.587 for *P. albocinereus* and *ρ* = 0.658 for *P. occidentalis*).

**Figure 3:**
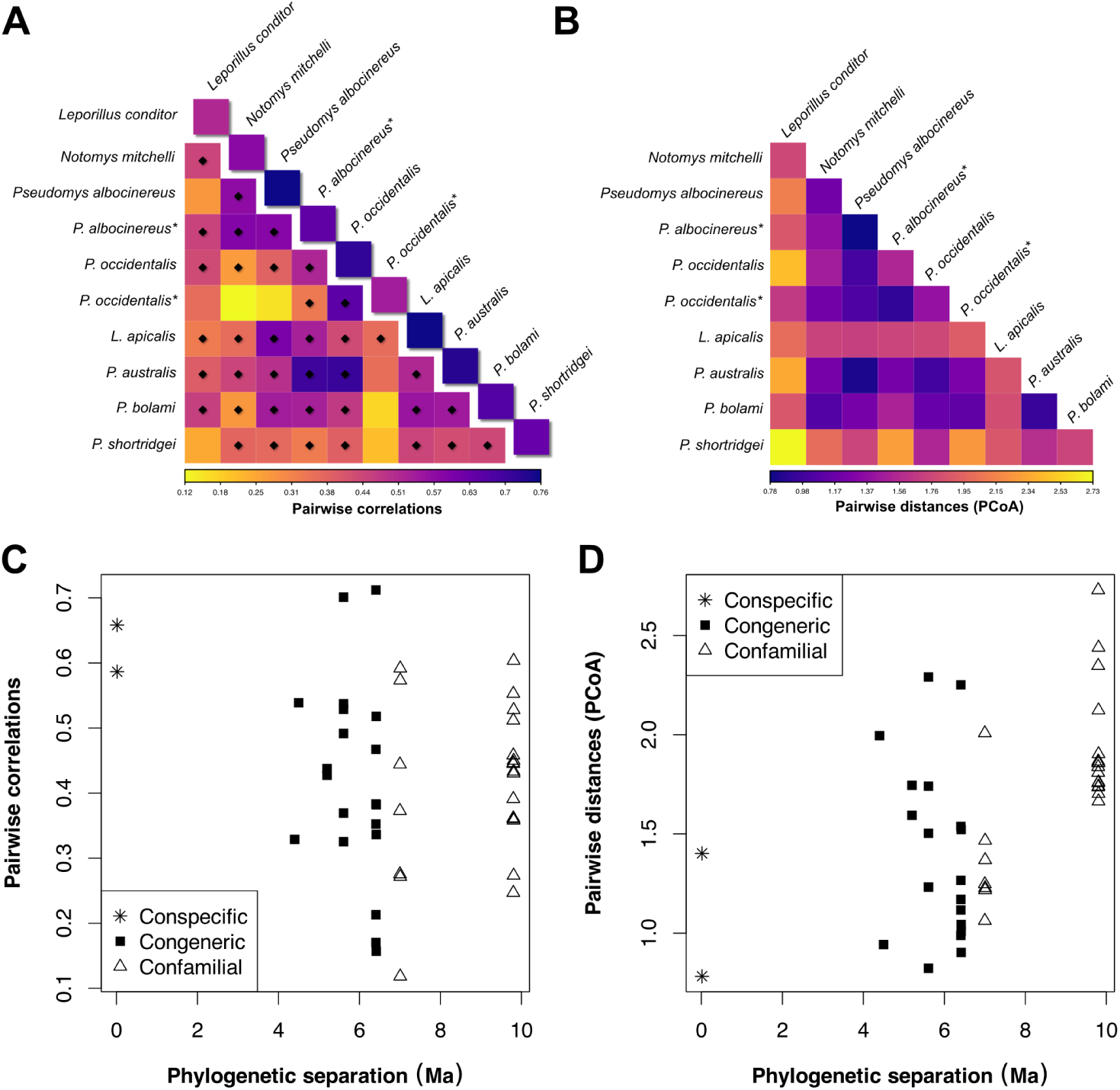
**A.** Pairwise correlations between structures of integration; correlations more similar than by chance by Mantel test indicated by diamonds; diagonal values are matrix repeatabilities; starred taxa are subsurface samples. **B.** Pairwise distances between samples in the Euclidean PCoA space resulting from relative eigenanalysis; colour scale reversed for more direct comparison with **A**. **C.** Pairwise correlations between structures of integration regressed against phylogenetic distance. **D.** Pairwise PCoA distances between samples regressed against phylogenetic distance. See supplement for tables of values.

### Relative eigenanalysis

Relative eigenanalysis indicates some discordance between phylogeny and the evolution of the structure of integration (Figure 3). Examination of the principal coordinates analysis does not show clear organisation by phylogenetic relationship and while the first principal coordinate axis might distinguish *Notomys* from other genera, higher axes reveal little phylogenetic organisation (Figure 4). Conspecifics are found generally near to one another in this space, but are in some cases closer to other samples (e.g. the subsurface samples of *Pseudomys occidentalis* and *P. albocinereus*).

**Figure 4:**
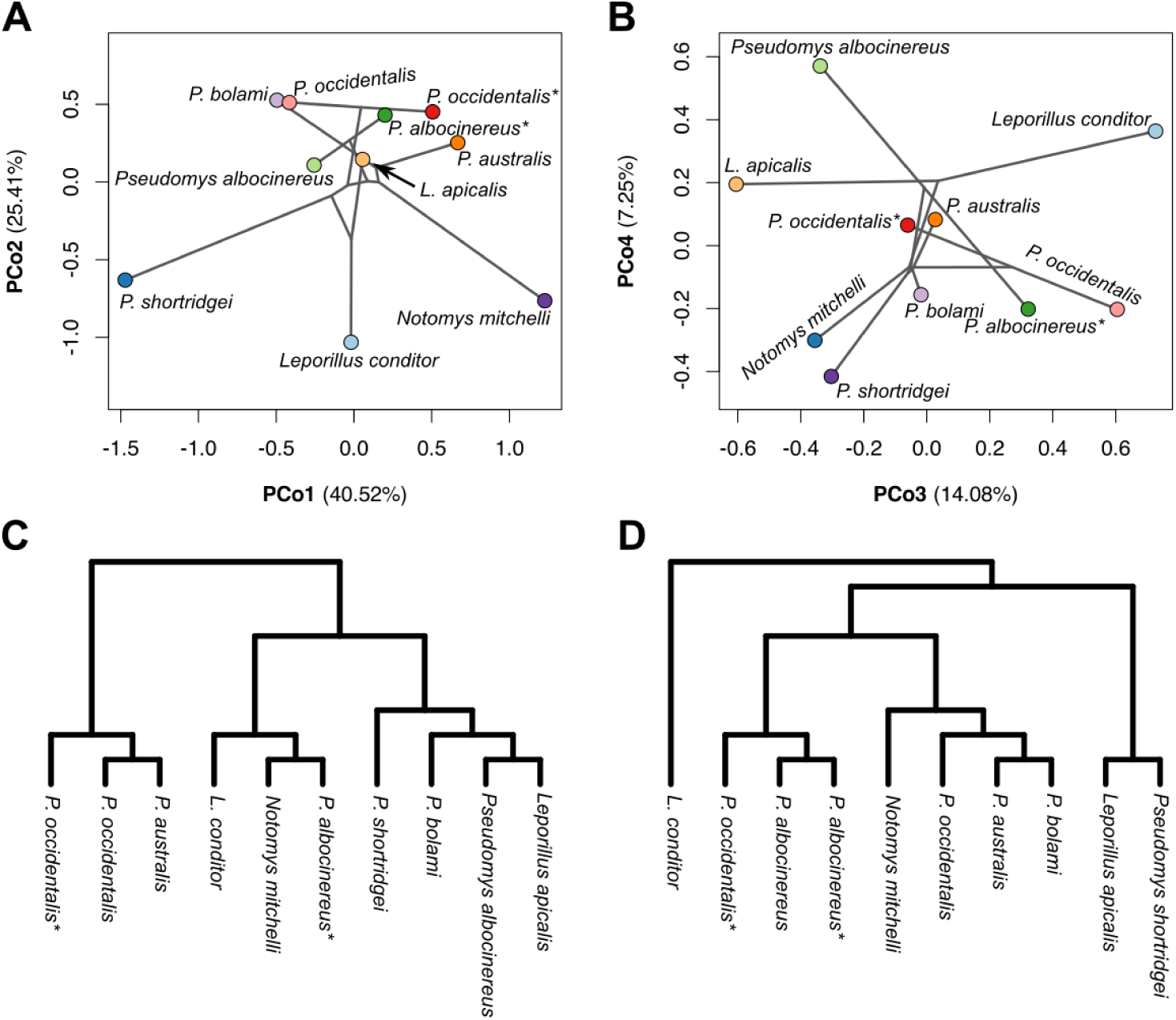
**A.** Relative eigenanalysis: PCo1 vs. PCo2; phylogeny from Figure 1A overlaid with node states estimated using the maximum likelihood fastAnc function in the R package phytools; starred taxa are subsurface samples. **B.** PCo3 vs. PCo4. **C.** Ward’s clustering of the integration disparity matrix; performed in the R package stats (version 4.4.2), see supplement for UPGMA clustering. **D.** Ward’s clustering of pairwise PCoA distances.

### Phylogenetic structure and the evolution of phenotypic integration

Clustering of the integration disparity matrix, either using Ward’s method (Figure 4C) or UPGMA (supplement), never resolves conspecifics closest to one another and clusters regularly show a closer affinity between confamilials than congenerics or even conspecifics. Species of the distantly related *Leporillus* are split among clusters and the topology of the cluster analyses is fundamentally inconsistent with that of the phylogeny. While this precise pattern is not mirrored in the equivalent analysis of the PCoA distance matrix (Figure 4D), similarity between some samples partially corresponds and the absence of other groupings persists (e.g. between species of *Leporillus*). Conspecifics of *Pseudomys albocinereus* are found to be most similar to one another in this case.

Regression of pairwise correlations between structures of integration against their corresponding phylogenetic distances reveals neither a uniform decay nor a consistent degree of similarity between structures of integration through time, at least in the range of 4 to 10 million years (Ma) of independent evolution (Figure 3C). Instead, the full range of observed similarities in structures of integration is recovered at about 6 Ma of phylogenetic separation. This range is less at about 10 Ma of independent evolution, but matches the central tendency (about *ρ* = 0.45) achieved by comparisons at some of the shortest evolutionary distances measured (4 to 5 Ma). A similar regression using pairwise PCoA distances may tell a slightly different story (Figure 3D): the spread of distances between 4 and 6.5 Ma of independent evolution includes comparisons with small values, but by 10 Ma of independent evolution, these are no longer present and the range has shifted to greater distances. While conspecific comparisons exhibit a fair degree of similarity in both the partition-based analyses and relative eigenanalysis, they are not the most similar comparisons using the former approach, and only the conspecific comparison of *Pseudomys albocinereus* is in the latter.

The integration disparity matrix contrasts with the phylogenetic relationships among the samples (Mantel test: *ρ* = 0.198, *p*-value = 0.149). A higher correspondence is found in the case of the PCoA distance matrix (*ρ* = 0.532, *p*-value = 0.012).

Comparison between the integration disparity matrix and the PCoA distance matrix reveals a weak correspondence between the two (Mantel test: *ρ* = 0.233, *p*-value = 0.153). These sets of pairwise comparisons are nonindependent from a phylogenetic standpoint but phylogenetic structure is not a strong predictor of either set of pairwise comparisons and correcting for relatedness only slightly weakens the correlation between the two matrices (supplement).

### Phylogenetic signal in trait associations

As expected, given the small number of taxa used here (10), only a few partial correlations between partitions are found to exhibit phylogenetic signal using either Pagel’s *λ* or Blomberg *et al*.’s *K* (Figure 2B, Figure 5). The partial correlations between the condyle and insertion of the superficial masseter, the ramus and anterior incisor alveolus, and between the posterior and anterior incisor alveolus all show significant phylogenetic signal, a result that is consistent between both metrics. Phylogenetic signal in the partial correlations between the condyle and proximal condyloid and between the distal condyloid and insertion of the internal pterygoideus are suggested by Pagel’s *λ* and found to be significant using Blomberg *et al*.’s *K*. Some other associations including those between the insertion of the superficial masseter and the proximal condyloid or between the molar and anterior incisor alveolus are more equivocally suggested to reflect phylogenetic signal (see supplement). Blomberg *et al*.’s *K* suggests variation in the partial correlations between the anterior incisor alveolus and ramus is particularly well explained by trait evolution under Brownian motion given the evolutionary relationships between the taxa investigated.

**Figure 5:**
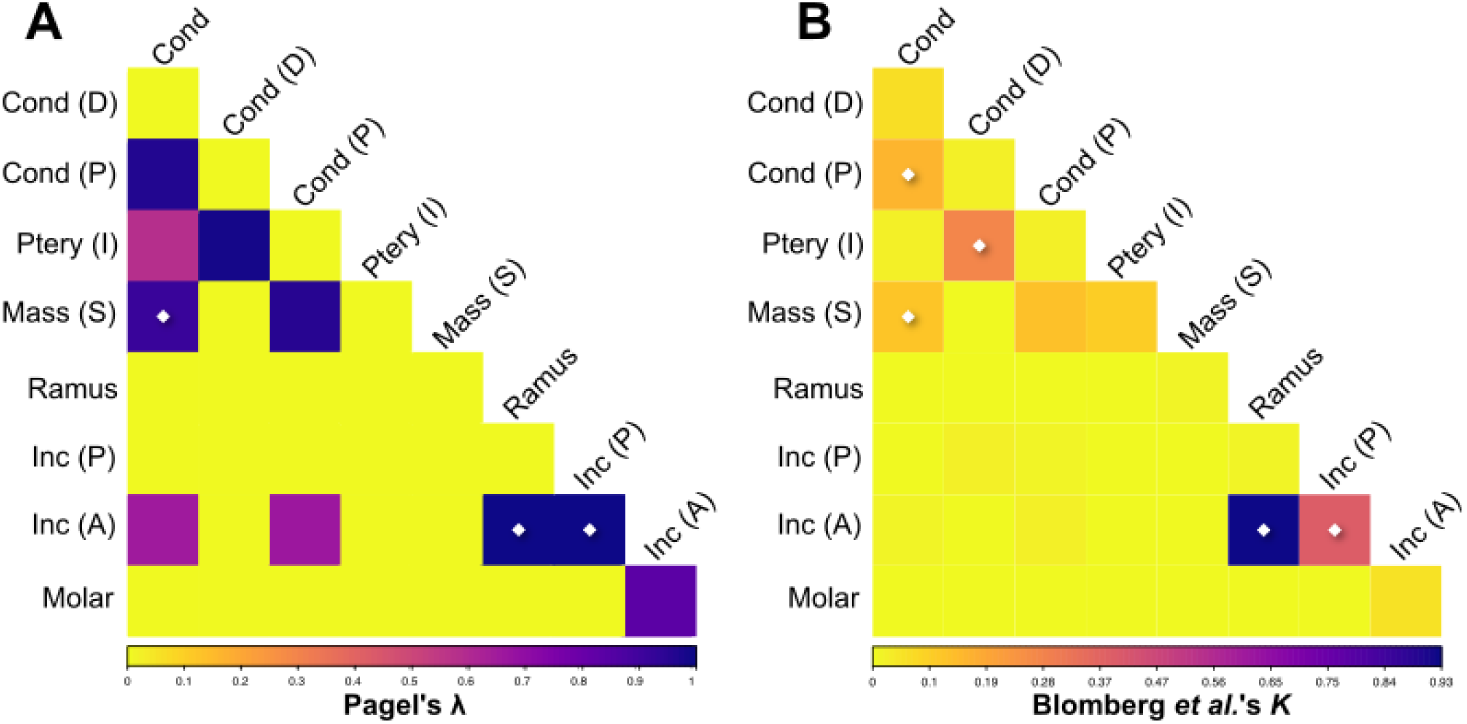
**A.** Pagel’s *λ* of partial correlations between partitions. **B.** Blomberg *et al*.’s *K* of partial correlations between partitions. Significant phylogenetic signal indicated by diamonds; tables of values in supplement. See Figure 1 for partition abbreviations.

## Discussion

The old endemic rodents are the tips of a radiation that may have begun as early as the late Miocene (Rowe *et al*., 2008). The results reported here show that mandibular integration is labile on timescales shorter than the radiation of this clade and that conservation and lability in different portions of the structure of integration have contributed to these patterns. This suggests that at least on the scale of this particular clade, the structure of integration as a whole could not have served as a strong constraint on the direction of phenotypic evolution. Still, on these timescales, the structure of integration is in most cases not subject to complete repatterning.

### Microevolution of the structure of integration

On the shortest timescales examined (thousands of years), the structure of integration was found to have evolved comparatively little (*Pseudomys albocinereus*: *ρ* = 0.587, *p*-value *<* 0.001; *P. occidentalis*: *ρ* = 0.658, *p*-value *<* 0.002). While most structures of integration are found to have diverged to a greater degree than this, not all have, for example in the comparison between the surface sample of *P. albocinereus* and *Leporillus apicalis* (*ρ* = 0.604, *p*-value *<* 0.001), species separated by about 10 Ma of independent evolution. Given the limited number of microevolutionary comparisons made here, it is difficult to broadly characterise how the structure of integration evolves on these timescales, but the possibility that it may undergo modification through the duration of a species cannot be precluded.

### Evolution of the structure of integration

None of the species sampled here share a common ancestor younger than about 2.2 million years before the present, and so it is not possible to characterise the initial pattern of conservation or decay among structures of integration. On longer timescales (about 4.4 Ma of independent evolution and greater), however, it is clear that the structure of integration has been subject to evolutionary modification. Unexpectedly, there is little definitive relationship between the duration of independent evolution (phylogenetic distance) and the degree to which two structures of integration have diverged (Figure 3C, D). Correlations between structures of integration are as low as 0.329 at about 4.4 Ma of independent evolution, but stronger correlations (*ρ >* 0.5) continue to be detected even at 10 Ma of phylogenetic separation. This range of correlations is consistent with a comparison between mandibular structures of integration from two rodent suborders (*ρ* = 0.482, *p*-value *<* 0.001, Zelditch *et al*., 2009). Despite the wide range of correlations, only in a few cases has the structure of integration been repatterned (e.g. in the comparison between the surface collection of *Pseudomys occidentalis* and *Notomys mitchelli*); most comparisons show significance for similarity, not dissimilarity. Relative eigenanalysis might detect a shift to more diverged structures of integration at about 10 Ma of independent evolution; this corresponds to comparisons made between the *Pseudomys* and *Conilurus* Divisions. Neither the correlations between structures of integration nor the pairwise distances derived from relative eigenanalysis yield a clear pattern among comparisons within a Division (corresponding to phylogenetic separations of less than about 6.5 Ma).

### Macroevolutionary patterns

Two major macroevolutionary patterns emerge. First, correlations between structures of integration show little relationship with phylogenetic distance on the scale of at least the Division, and possibly the focal clade at large: comparisons between structures of integration produce both low and high correlations at all sampled macroevolutionary scales (Figure 3C). Second, despite the wide range of correlation strengths, most comparisons between structures of integration are shown to be more similar than by chance (Figure 3A).

While it is not surprising that the degree of similarity between structures of integration will erode before statistical similarity is lost, these results indicate that a uniform decrease of similarity is not the dominant pattern of evolution on the observed timescales. The detection of high correlations between structures of integration in long-diverged species (Figure 3C), the absence of an obvious phylogenetic pattern of diffusion through PCoA space (Figure 4A, B), and the poor correspondence between phylogenetic structure and the pattern of correlations between structures of integration all strongly suggest this. At timescales longer than those investigated here it seems inevitable that statistical similarity would be lost between structures of integration but, prior to this, the structure of integration is already evolving and apparently not in ways that trend solely towards dissimilarity.

It is difficult to say why this might be the case or how lability in a complex structure of integration might lend itself to apparent convergence as much as it does divergence, but correlational selection might play a role (Lande and Arnold, 1983; Svensson *et al*., 2021) and analogous comparisons of cranial structures of integration in primates show this pattern is not without precedent (Mitteroecker and Bookstein, 2008; Ackermann, 2009). These unintuitive findings are perhaps symptomatic of a common challenge to the interpretation of patterns of phenotypic integration. These patterns are readily analysed at a particular focal level, and yet contributions to phenotypic integration are distributed across a variety of levels of biological organisation (genome, epigenome, phenotype, ontogeny, etc.) producing a palimpsest-like developmental architecture (Hallgrímsson *et al*., 2006; Hallgrímsson *et al*., 2009) that may evolve in unpredictable ways (e.g. Jamniczky and Hallgrímsson, 2011). Because of this, a closer examination of specific trait associations can offer hints at possible drivers to the observed patterns. Ultimately, while the wide range of strengths of correlations between structures of integration is clear evidence that the structure itself is evolving, the strength of a correlation alone says very little about which portions of the structure of integration contribute to similarities and differences in the final metric.

### Conservation and lability within the structure of integration

The structures of integration measured here record the conservation of only a few strong trait associations (Figure 2A). This suggests that the partition boundaries defined in this study routinely divide these conserved modules. These cases of integration all involve topologically adjacent partitions, consistent with “neighbourhood effects” observed in other studies (Chernoff and Magwene, 1999; Klingenberg and Zaklan, 2000; Klingenberg *et al*., 2001; Zelditch *et al*., 2009; Webster and Zelditch, 2011a) and suggestive of local morphogenetic control. While other non-adjacent partitions are revealed to be phenotypically integrated, these associations are not well conserved between the structures studied (interestingly a similar pattern was found among three species of Cambrian trilobites [Webster and Zelditch, 2011a]). At evolutionary scales, similarity between structures of integration may be driven by the persistence of conserved neighbourhood effects while lability in delocalised trait associations may contribute to dissimilarity. Concordant evolution within these localised modules but not between them may also drive net dissimilarity between structures of integration.

Lability in some trait associations shows measurable phylogenetic signal (Figure 2B, Figure 5) and evolution of function may play a role here. As discussed by Zelditch *et al*. (2009), the material characteristics of diet contribute to an individual’s reliance either on gnawing or chewing behaviours and this might influence the position of muscle attachment (see also Menegaz and Ravosa, 2017). Phylogenetic associations including the condyle, insertion of the internal pterygoideus, and insertion of the superficial masseter may be evidence of such a dietary/behavioural signal as witnessed by the coordinated evolution of the articulating geometry of the condyle and the shape of attachment of these two muscles. A complimentary interpretation might be proposed for the phylogenetic association involving portions of the incisor alveolus and ramus. Interestingly, the mammalian mandible has often been divided into an anterior tooth-bearing module and a posterior module defined by articulation and muscle attachment (Klingenberg *et al*., 2003). This two-module scheme is broadly consistent with the two sets of trait associations identified above as expressing phylogenetic signal. Nevertheless, it is important to emphasise that phylogenetic signal in the association between traits does not demonstrate the detection of a conserved variational module (as is accomplished in direct comparisons between correlation structures), nor does it necessarily reflect a pattern of evolutionary integration (which is not explored in this study). Rather, it shows that variation between taxa, in some aspect of their structures of integration, is well described by a model of trait evolution (in this case Brownian motion) and the evolutionary relationships between the taxa.

### Possible drivers and future work

In a clade which has radiated into a range of dietary niches (such as the old endemic rodents of Sahul), it is conceivable that shifts in functional demands, mediated by selection, might induce repatterning in the structure of integration. A similar function-oriented argument has been put forth for evolution of both mandibular (Monteiro and Nogueira, 2010) and cranial (Rossoni *et al*., 2019) integration in phyllostomid bats, and in evolution of cranial integration in a genus of vespertilionid bat (Dzeverin and Vertsimakha, 2024). A radiation characterised by functional adaptation might reshuffle the structure of integration as certain trait associations are indirectly drawn across an adaptive landscape by selection.

A more complex model involving phylogeny, diet, and estimation of force regimes across the mandible (e.g. using finite element analysis [Tsouknidas *et al*., 2017] or including patterns of geographic variation in tooth wear [Schram and Turnbull, 1970]) might more clearly describe the link between particular trait associations and the evolution of the structure of integration. At the same time, given statistical limitations on estimating phylogenetic signal, a larger sample of structures of integration would permit a close examination of phylogenetic signal across the evolution of the constituent trait associations and could potentially illuminate more developmentally localised effects of selection, developmental constraint, and stasis. Broad phylogenetic sampling would also allow for more sophisticated investigation of the evolution of trait associations under a variety of models of trait evolution; labile elements in the structure of integration should not be expected to all evolve under the same dynamics.

Even a few estimates of the structure of phenotypic integration can provide substantial inferential power. A more robust and nuanced predictive framework for developmental bias on phenotypic evolution might involve the subset of conserved elements of the structure of integration, derived from even a few taxa. Precisely the same framework will also suggest traits and trait associations that are expected to be less constrained in their directions of phenotypic evolution due to their more labile developmental underpinnings.

The present study is unable to address several important questions that future work should seek to investigate. First, with the exception of two intraspecific comparisons, it works in essentially ultrametric terms with little access to ancient phenotypes close to nodes in the phylogeny. The fossil record provides a rich array of phenotypes that can be accessed through the evolutionary history of clades and a careful study of the evolution of phenotypic integration can yield valuable information on the precise dynamics of the evolution of development (Hunt, 2007; Webster and Zelditch, 2011a). Second, because few intraspecific comparisons are made here, it is difficult to more broadly characterise microevolutionary patterns with regards to development. Where due attention is paid to the effects of taphonomy and time-averaging, the fossil record is once again a natural place to examine development through the duration of a species (e.g. Goswami *et al*., 2015). Cave deposits (like those examined here) provide rich microevolutionary records and many have abundant material capturing species ranges which span important recent environmental fluctuations (Rowe and Terry, 2014; Hunt, 2004). Numerous planktonic organisms leave an exquisite microevolutionary fossil record which can be accessed in outcrop or by drill core (Hunt, 2007). A parallel line of inquiry which does not require access to deep time is to investigate whether geographic variation has any bearing on the structure of integration, although expectations remain unclear (compare Malvolti *et al*., 1994; Roff *et al*., 2004; Swiderski and Zelditch, 2022). As O’Meara and Beaulieu (2024) note, measurement error, especially at the microevolutionary scale, must be carefully considered. Finally, broader phylogenetic sampling appears necessary to capture the complete loss of similarity between structures of integration. This scale might be broader than clear homology will permit: divergence among, but overall conservation of, cranial structures of integration has been reported in catarrhine primates (de Oliveira *et al*., 2009), phyllostomid bats (Rossoni *et al*., 2019), ruminants (Haber, 2015), carnivorans (Machado *et al*., 2018), and even therian mammals (Goswami, 2006; Porto *et al*., 2009). Dense phylogenetic sampling will also permit a robust characterisation of the evolutionary structure of integration from which valuable comparisons can be made with intraspecific structures of integration to assess patterns of developmental constraint.

## Conclusion

The relationship between the developmentally mediated production of variation and the evolutionary sorting of that variation is a critical component to many emerging perspectives on evolutionary biology but remains incompletely understood (Webster, 2019; Jablonski, 2020; Holstad *et al*., 2024; Stansfield and Parsons, 2024). The work presented here offers some hints at the lability of phenotypic integration on macroevolutionary timescales and the forces that may propel evolutionary modification to development, but it is clear that further investigation on almost every scale of biological organisation will be necessary to fully appreciate the influence of development on phenotypic evolution and the malleability of developmental systems in the face of selection.

## Notes

### Competing Interest Statement

The authors have declared no competing interest.

