## Supplement for "Conservation and lability within the structure of mandibular integration in “old endemic” Australian rodents": Distance_matrix.R

```
#### Distance matrix and correlation structure calculations

# Preamble
  library(geomorph)
  library(shapes)
  library(vegan)

# Landmark data as an n x p x k (landmarks, dimensions, specimens) array.
  # These example data (surface collection of Pseudomys albocinereus) have had semi-landmarks slid to minimise bending energy and the number of semi-landmarks reduced.
    lm_data <- readland.tps("Example_LMs_for_script.tps", specID = "ID")

# List of partitions, each named element is a vector of landmarks to include in a partition
  part_list <- list(cond = c(5:7),
                    cond_d = c(22:25), 
                    cond_p = c(18:21),
                    ptery = c(8, 12:17),
                    mass = c(28:39),
                    ramus = c(44:47, 9, 26:27),
                    inc_p = c(10, 40:43, 11, 52:55),
                    inc_a = c(1, 2, 56:59),
                    molar = c(3,4, 48:51))

# Function to generate matrix correlations between partitions, takes n x p x k landmark array and named list of partitions (as above).  1,000 permutation Mantel test used.
  cor_struct_gen <- function(lm_data, part_list){
    n_parts <- length(part_list)
    # Superimpose individual partitions separately
      sup_parts <- list()
      for(p in 1:n_parts){
        sup_parts[[p]] <- gpagen(lm_data[part_list[[p]],,])$coords
      }


    # Pairwise Procrustes distance of a partition between each individual (distance matrix for each partition)
      dist_parts <- list()
      for(p in 1:n_parts){
        samp_size <- dim(lm_data)[3]
        dist_mat <- matrix(nrow = samp_size, ncol = samp_size)
        for(n in 1:(samp_size-1)){
          for(q in (n+1):samp_size){
            dist_mat[q,n] <- procdist(x = sup_parts[[p]][,,n], y = sup_parts[[p]][,,q])
          }
        }
        dist_parts[[p]] <- dist_mat
      }

    # Matrix correlation between distance matrices
      cor_mat <- matrix(nrow = n_parts, ncol = n_parts)
      dimnames(cor_mat) <- list(names(part_list), names(part_list))
      cor_mat_p <- cor_mat
      for(n in 1:(n_parts-1)){
        for(p in (n+1):n_parts){
          temp_mantel <- mantel(xdis = dist_parts[[n]], ydis = dist_parts[[p]], permutation = 1000)
          cor_mat[p,n] <- temp_mantel$statistic
          cor_mat_p[p,n] <- temp_mantel$signif
        }
      }

    # Return correlation structure and significance values as distance matrices
      results <- list("Mantel matrix corr." = cor_mat,
                      "P-values" = cor_mat_p)
      return(results)
  }

# Run script
  # Compare example result to Supplement, Table 3 (p-values will vary slightly due to Mantel permutations).
    cor_struct_P_albocinereus <- cor_struct_gen(lm_data, part_list)
```
