## Supplement for "Conservation and lability within the structure of mandibular integration in “old endemic” Australian rodents": Supplement.pdf

**4 Exploring the structure of integration within a sample** **6**

**5 Clustering of the integration disparity matrix and matrix of PCoA distances** **6**

**6 Supplemental Tables** **7**

**7 Supplemental Figures** **13**

**References** **31**

### 1 Description of supplemental files

- `RawLMdata.tps`: raw landmark data, scale of coordinates is 1 millimetre, landmark configurations associated with specimen IDs. Includes semi-landmarks before reduction in number.

- `Distance_matrix.R`: script for generating a distance matrix between partitions of landmark data and calculating the correlation structure.

- `Example_LMs_for_script.tps`: example prepared landmark data (surface sample of *Pseudomys albocinereus*) for use in `Distance_matrix.R` script.

- `Specimens.csv`: specimen information including genus, species, ID, side, size (length measured between landmarks 2 and 6), locality, horizon, type of sample (surface vs. subsurface). Locality abbreviations followed by sampling horizon in inches below surface: WC = Weeke's Cave, HC = Hastings Cave, MEEC = Murra-el-elevyn Cave.

### 2 Sample background

#### 2.1 Previous work

The "old endemic" (Simpson, 1961) rodents of Sahul (Australia + New Guinea) represent a radiation from southeast Asia that likely began in the late Miocene or early Pliocene (Rowe *et al.*, 2008). This radiation is characterised by a flourishing of ecomorphotypes, body sizes, diet, locomotory strategies, and modes of sociality with many cases of phenotypic convergence that echo other major rodent radiations. This radiation took place through a period of substantial environmental change and variability (Fujioka and Chappell, 2010) and the fossil record of even extant species shows how geographic ranges have shifted and in some cases become disjunct since the Pleistocene (Lundelius, 1957). One primarily Australian group of old endemic rodents is the traditionally designated "Conilurini". Molecular phylogenetics have since demonstrated another tribe, the Uromyini to be nested within the "Conilurini", sister to what is now known as the *Conilurus* Division and to the exclusion of the *Pseudomys* Division (Rowe *et al.*, 2008; Roycroft *et al.*, 2020, 2021). The species studied here are from the *Conilurus* and *Pseudomys* Divisions (Figure 1).

During the mid to late 20$^{th}$ Century, major collections of subfossil vertebrate material were made from a number of caves in South and Western Australia (Figure 1). Collections from Madura Cave in the Nullarbor Plain (Western Australia) were the primary focus of the comprehensive reports that emerged from this large sampling effort (Lundelius and Turnbull, 1973, 1975, 1978, 1981, 1982, 1984, 1989, 1999). Many other caves were also extensively sampled (Lundelius, 1957, 1960, 1963, 1983) and much of the resulting material would ultimately be housed in the collections of the Field Museum of Natural History, Chicago, USA. These substantial collections have received only minimal recent attention (e.g. Koeller and Mitchell, 2014), but they provide ample opportunity to investigate a variety of important questions including those pertaining to the role of phenotypic integration on evolutionary disparification.

A robust assessment of the structure of phenotypic integration should be made at the population level, which is most reliably accomplished in live populations. However cave deposits offer several distinct advantages. These assemblages can provide large sample sizes for elusive, endangered, or even extinct taxa (Lundelius, 1983; Vakil *et al.*, 2023) in regions where extended field efforts are challenging. The stratified nature of cave deposits also permits a sequence of samples documenting the microevolutionary history of a species, a type of sampling unavailable to even long-term live-only studies.

#### 2.2 Localities and sampling

Hastings Cave is in Drovers Cave National Park, near Jurien Bay in Western Australia (Figure 1). Like Madura Cave, Hastings Cave was subjected to stratified sampling but unlike Madura Cave the lowest sampled layers are much younger; post-Pleistocene (Lundelius, 1983), and perhaps on the order of 10,000 years before present (Y.B.P.). The preservation at Hastings Cave is generally superior to that at Madura Cave with numerous intact hemimandibles of *Pseudomys albocinereus* and *P. occidentalis.* These species have a lower set of occurrences, older than 7,850 $\pm$ 170 Y.B.P. (Lundelius, 1960, Hastings Cave therein referred to as Drover's Cave), and a younger surface sample, allowing for characterisation of microevolutionary effects on patterns of development (see also Lundelius, 1983). For both species, the lower set of occurrences was treated as one sample with the surface collections treated as a second sample. *Pseudomys shortridgei* is also found at Hastings Cave, but in sufficient abundance only to derive a sample from the surface collections.

Murra-el-elevyn Cave in the Nuytsland Nature Reserve of the Nullarbor Plain in southeast Western Australia (Figure 1) is represented by a rich surface collection of small mammal skeletal material. From Murra-el-elevyn Cave, the recently extinct (Woinarski and Burbidge, 2016) *Leporillus apicalis* was sampled alongside the extant *Pseudomys australis* and *P. bolami.* The age of a surface collected mummified *Thylacinus cynocephalus* has been radiometrically dated at 3,280 $\pm$ 90 Y.B.P. (Partridge, 1967), however this specimen was discovered deeper into the cave and is probably not representative of the collections (Lundelius, 1963) studied here.

At Weekes Cave in the Nullarbor Wilderness Protection Area, southwestern South Australia (Figure 1), surface collections provide a rich sampling of well-preserved hemimandibles of recent *Notomys mitchelli* and *Leporillus conditor.* As with the specimens from Murra-el-elevyn Cave, clear radiometric dates are not available (for comments on these deposits see Baird, 1990), but this collection is probably restricted to a recent interval of time given the surface sampling effort and as suggested by the lack of obvious staining and biostratinomic wear of the material compared with the style of preservation in older and deeper horizons at other caves.

#### 2.3 Geographic and temporal variation

Many of the taxa investigated here range (or historically ranged) through substantial regions of south central and southwestern Australia. Although attenuation of gene flow is weakly detectable only at the continental scale in widely distributed species (e.g. Roycroft *et al.*, 2021), geographic variation has been documented in the morphology of some of the species studied here (Lundelius, 1964). Nevertheless, the effects of geographic variation are deemed minimal given that each species is sampled from a single cave and each cave assemblage is restricted to rodent populations within the hunting ranges of local aerial predators. These hunting ranges are dwarfed by the biogeographic ranges of prey species (Marshall, 1986; Lundelius, 1957).

The cave deposits studied here are primarily accumulations of disaggregated owl pellets which have amassed on timescales ranging from historic to late Pleistocene and are subject to time-averaging, temporal mixing which can result in the co-occurrence in the same sample of individuals that lived at different times (Kowalewski, 1996). Near-surface cave assemblages have been shown to have high fidelity when compared to the composition of living faunas that continue to contribute to them (Terry and Novak, 2015). At a few of the Australian caves, excavations permit examination of a deeper, stratified sequence of intervals. Without a flux of recent material into these assemblages, the temporal structure of these deeper samples is expected to be more influenced by time-averaging (Tomašovỳch *et al.*, 2023), nevertheless, the effects of time-averaging in sub-surface cave deposit samples have shown mammalian dental trait variances and covariances to be fairly good approximations of population-level values (Hunt, 2004a). Analytical time-averaging (the pooling of data from multiple stratigraphic samples) also seems to only slightly inflate trait variance with similar effects expected in trait covariance (Hunt, 2004b, p. 503). These results suggest relative stability of these properties on within-lineage timescales of evolution, although how this phenomenon manifests in more sensitive analyses of complex structures of phenotypic integration remains to be seen.

### 3 Specimens and shape analysis

#### 3.1 Landmark selection

The coronoid process, anterior margin of the condylar process, and ventral margin of the incisor alveolus are taphonomically vulnerable, and were excluded from the analysis so as to increase sample size. Landmark selection permitted inclusion of edentulous hemimandibles.

#### 3.2 Landmark descriptions

1. Point at which upper surface of incisor enters the incisor alveolus, fairly visible even when incisor is missing.

2. Dorsalmost extent of incisor alveolus.

3. Point of inflection between posteriorly rising limb of diastema and position of M1.

4. Point of posteriormost extent of tooth row and base of coronoid process.

5. Anterior boundary between condylar surface and condyloid process.

6. Posterior boundary between condylar surface and condyloid process.

7. Point of inflection between posterodorsal curvature of condyloid process and condyloid.

8. Tip of angular process.
9. Dorsalmost point of ventral curvature along ramus, typically associated with masseteric arch.
10. Dorsalmost point of ventral curvature along incisor alveolus, typically delineated by a shallow reinforced ridge or "chin".
11. Position of mental foramen.

Semi-landmark curves are between landmarks 2 and 3, 7 and 8, 8 and 9, and 9 and 10.

#### 3.3 Left vs. right hemimandibles

Left and right hemimandibles from the same individual should not be treated as independent. However, the nature of preservation in these collections means that hemimandibles are almost exclusively found in isolation. In cases where sample size permitted, hemimandibles from the same side were used to avoid this problem. In some cases, an analytically robust sample size could only be achieved by including hemimandibles of both sides. Low sample size in these cases is not due to few preserved individuals (which would imply a high likelihood of sampling both left and right hemimandibles from one individual), but rather is the result of pronounced biostratinomic degradation which rendered most hemimandibles unviable for landmarking. Indeed, it is for this reason that the best studied cave by Lundelius and Turnbull (Madura Cave) could not be used in the present study: too few hemimandibles were of sufficient preservational quality to adequately support geometric morphometric analysis.

#### 3.4 Measurement error

To assess error in photographic orientation, a single hemimandible (*Pseudomys albocinereus*, surface sample) was independently mounted and photographed and digitised 20 times over the course of about four months. Digitisation error was quantified through independent landmarking of a single image of the aforementioned *P. albocinereus* hemimandible 20 times over the same timeframe. Procrustes variances were calculated for each set of landmark configurations.

Procrustes variances show these sources of error are between one and two orders of magnitude smaller than within-sample variation: mounting error = 0.0000578, digitising error = 0.0000420, surface sample of *Pseudomys albocinereus* = 0.000945. A morphospace of these data similarly shows this (Figure 2).

#### 3.5 Correcting for allometry

A linear regression of shape variables on the natural logarithm of centroid size (lnCS) allows shape to be predicted at a specified size (here mean lnCS) and residuals (shape deviations) from the regression, associated with each specimen, are added to this predicted shape, effectively removing size-related shape variation from the data (Zelditch *et al.*, 2012). Allometry and associated statistics (using the adonis2 function in the R package `vegan` [version 2.6.10]) are shown in Figure 3 and Figure 4.

### 4 Exploring the structure of integration within a sample

Three exploratory methods were used in this study: clustering, reticulate network analysis, and graphical modelling. These methods have been used to analyse correlation matrices in previous studies (Monteiro *et al.*, 2005; Zelditch *et al.*, 2008, 2009; Webster and Zelditch, 2011b,a) and details on those methods can be found there. These methods were implemented in largely the same fashion as they have been previously with more recent software packages used in a few cases:

- Clustering (Ward's and UPGMA): run in R (version 4.4.2) using the `stats` package (version 4.4.2). Cophenetic correlations calculated in the aforementioned package. Agglomerative coefficients calculated in the `cluster` package (version 2.1.6).
- Reticulate network analysis: run on the T-REX web server (Boc *et al.*, 2012, `http://www.trex.uqam.ca/`) on 17 January 2025 using the "ADDTREE" option and conservative Q1 criterion.
- Graphical modelling: run in R using the `gRim` package (Højsgaard *et al.*, 2012, version 0.3.4) using both "headlong" and "all" search options; results corresponded to those from the commonly used MIM (version 3.2.0.7).

See supplemental figures (Figure 5 through Figure 15) and tables (Table 1 through Table 10) below for correlation matrices between partitions and results of exploratory analyses.

### 5 Clustering of the integration disparity matrix and matrix of PCoA distances

UPGMA clustering of the integration disparity matrix and pairwise PCoA distances is shown in Figure 19. Clustering statistics for both Ward's and UPGMA clustering can be found in Table 17.

### 6 Supplemental Tables

| | Cond | Cond (D) | Cond (P) | Ptery (I) | Mass (S) | Ramus | Inc (P) | Inc (A) | Molar |
|---|---|---|---|---|---|---|---|---|---|
| Cond | | **0.039** | 0.302 | 0.642 | **0.040** | **0.009** | **0.015** | 0.079 | 0.702 |
| Cond (D) | 0.142 | | **0.042** | 0.808 | 0.668 | 0.059 | **0.042** | 0.309 | 0.749 |
| Cond (P) | 0.029 | 0.197 | | 0.343 | 0.580 | 0.446 | 0.199 | 0.694 | 0.526 |
| Ptery (I) | <0.001 | <0.001 | 0.041 | | **0.021** | 0.073 | 0.059 | 0.512 | 0.496 |
| Mass (S) | 0.134 | <0.001 | <0.001 | 0.221 | | 0.107 | 0.220 | 0.449 | 0.451 |
| Ramus | 0.206 | 0.164 | 0.012 | 0.159 | 0.113 | | **<0.001** | 0.319 | 0.375 |
| Inc (P) | 0.175 | 0.211 | 0.096 | 0.201 | 0.070 | 0.406 | | 0.102 | 0.912 |
| Inc (A) | 0.134 | 0.052 | <0.001 | <0.001 | 0.005 | 0.044 | 0.166 | | 0.944 |
| Molar | <0.001 | <0.001 | <0.001 | <0.001 | 0.009 | 0.023 | <0.001 | <0.001 | |

**Table 1:** Correlation structure (lower triangle) of *Leporillus conditor*; Mantel test $p$-values (upper triangle).

| | Cond | Cond (D) | Cond (P) | Ptery (I) | Mass (S) | Ramus | Inc (P) | Inc (A) | Molar |
|---|---|---|---|---|---|---|---|---|---|
| Cond | | **0.015** | 0.121 | 0.157 | **0.036** | 0.880 | 0.498 | 0.140 | 0.070 |
| Cond (D) | 0.155 | | 0.200 | 0.590 | 0.372 | 0.189 | 0.620 | 0.071 | 0.553 |
| Cond (P) | 0.074 | 0.064 | | 0.250 | 0.906 | 0.192 | 0.100 | 0.101 | 0.139 |
| Ptery (I) | 0.070 | <0.001 | 0.045 | | **<0.001** | 0.579 | 0.134 | 0.253 | 0.081 |
| Mass (S) | 0.105 | 0.017 | <0.001 | 0.323 | | **0.041** | 0.297 | 0.272 | 0.190 |
| Ramus | <0.001 | 0.065 | 0.061 | <0.001 | 0.126 | | **<0.001** | 0.111 | 0.650 |
| Inc (P) | <0.001 | <0.001 | 0.120 | 0.102 | 0.034 | 0.365 | | **0.025** | 0.185 |
| Inc (A) | 0.078 | 0.117 | 0.109 | 0.048 | 0.037 | 0.102 | 0.221 | | 0.549 |
| Molar | 0.103 | <0.001 | 0.080 | 0.113 | 0.061 | <0.001 | 0.074 | <0.001 | |

**Table 2:** Correlation structure (lower triangle) of *Notomys mitchelli*; Mantel test $p$-values (upper triangle).

| | Cond | Cond (D) | Cond (P) | Ptery (I) | Mass (S) | Ramus | Inc (P) | Inc (A) | Molar |
|---|---|---|---|---|---|---|---|---|---|
| Cond | | **0.006** | 0.946 | 0.441 | 0.703 | 0.696 | 0.396 | 0.219 | 0.083 |
| Cond (D) | 0.221 | | **0.048** | 0.481 | 0.805 | 0.251 | 0.715 | 0.225 | 0.801 |
| Cond (P) | <0.001 | 0.138 | | **<0.001** | **0.017** | 0.648 | 0.475 | 0.817 | 0.201 |
| Ptery (I) | 0.004 | <0.001 | 0.322 | | **<0.001** | 0.495 | 0.414 | 0.155 | **0.013** |
| Mass (S) | <0.001 | <0.001 | 0.193 | 0.448 | | **0.002** | 0.141 | 0.467 | **0.009** |
| Ramus | <0.001 | 0.030 | <0.001 | <0.001 | 0.218 | | **<0.001** | 0.568 | 0.169 |
| Inc (P) | 0.010 | <0.001 | <0.001 | 0.008 | 0.078 | 0.417 | | **0.048** | **0.014** |
| Inc (A) | 0.045 | 0.045 | <0.001 | 0.058 | 0.003 | <0.001 | 0.106 | | **0.011** |
| Molar | 0.093 | <0.001 | 0.054 | 0.166 | 0.189 | 0.049 | 0.177 | 0.152 | |

**Table 3:** Correlation structure (lower triangle) of *Pseudomys albocinereus*; Mantel test $p$-values (upper triangle).

| | Cond | Cond (D) | Cond (P) | Ptery (I) | Mass (S) | Ramus | Inc (P) | Inc (A) | Molar |
|---|---|---|---|---|---|---|---|---|---|
| Cond | | **0.007** | 0.879 | 0.721 | 0.527 | 0.625 | 0.621 | 0.645 | 0.917 |
| Cond (D) | 0.265 | | 0.154 | 0.375 | 0.078 | 0.221 | 0.507 | 0.636 | 0.473 |
| Cond (P) | <0.001 | 0.136 | | 0.349 | 0.288 | 0.469 | 0.397 | 0.081 | 0.154 |
| Ptery (I) | <0.001 | 0.015 | 0.032 | | **0.006** | **0.044** | 0.234 | 0.783 | 0.351 |
| Mass (S) | <0.001 | 0.196 | 0.060 | 0.300 | | **0.006** | 0.066 | 0.303 | 0.587 |
| Ramus | <0.001 | 0.035 | <0.001 | 0.229 | 0.362 | | **<0.001** | 0.435 | 0.623 |
| Inc (P) | <0.001 | <0.001 | 0.020 | 0.066 | 0.225 | 0.710 | | 0.080 | 0.267 |
| Inc (A) | <0.001 | <0.001 | 0.168 | <0.001 | 0.068 | <0.001 | 0.205 | | 0.167 |
| Molar | <0.001 | <0.001 | 0.131 | 0.023 | <0.001 | <0.001 | 0.056 | 0.141 | |

**Table 4:** Correlation structure (lower triangle) of *Pseudomys albocinereus* (subsurface); Mantel test $p$-values (upper triangle).

| | Cond | Cond (D) | Cond (P) | Ptery (I) | Mass (S) | Ramus | Inc (P) | Inc (A) | Molar |
|---|---|---|---|---|---|---|---|---|---|
| Cond | | **<0.001** | 0.380 | 0.601 | 0.079 | **0.022** | **0.044** | 0.841 | 0.069 |
| Cond (D) | 0.399 | | 0.724 | 0.532 | 0.063 | 0.064 | **0.021** | 0.829 | **0.020** |
| Cond (P) | 0.020 | <0.001 | | **0.002** | 0.077 | **0.045** | 0.062 | 0.437 | 0.892 |
| Ptery (I) | <0.001 | <0.001 | 0.278 | | 0.337 | 0.112 | 0.313 | 0.200 | 0.561 |
| Mass (S) | 0.150 | 0.122 | 0.124 | 0.029 | | 0.095 | 0.072 | 0.917 | 0.901 |
| Ramus | 0.191 | 0.111 | 0.139 | 0.096 | 0.106 | | **<0.001** | 0.619 | 0.104 |
| Inc (P) | 0.162 | 0.159 | 0.139 | 0.038 | 0.136 | 0.599 | | 0.104 | **0.002** |
| Inc (A) | <0.001 | <0.001 | 0.015 | 0.061 | <0.001 | <0.001 | 0.103 | | 0.387 |
| Molar | 0.110 | 0.146 | <0.001 | <0.001 | <0.001 | 0.086 | 0.197 | 0.020 | |

**Table 5:** Correlation structure (lower triangle) of *Pseudomys occidentalis*; Mantel test $p$-values (upper triangle).

| | Cond | Cond (D) | Cond (P) | Ptery (I) | Mass (S) | Ramus | Inc (P) | Inc (A) | Molar |
|---|---|---|---|---|---|---|---|---|---|
| Cond | | **0.004** | 0.252 | 0.548 | 0.229 | **0.008** | **0.017** | 0.598 | 0.108 |
| Cond (D) | 0.325 | | 0.743 | 0.254 | 0.127 | 0.068 | 0.307 | 0.256 | 0.075 |
| Cond (P) | 0.062 | <0.001 | | 0.151 | 0.474 | 0.087 | 0.644 | 0.730 | 0.286 |
| Ptery (I) | <0.001 | 0.082 | 0.104 | | 0.428 | 0.065 | 0.303 | 0.843 | **0.050** |
| Mass (S) | 0.090 | 0.160 | 0.005 | 0.026 | | 0.247 | 0.102 | 0.274 | 0.415 |
| Ramus | 0.295 | 0.223 | 0.145 | 0.185 | 0.092 | | **<0.001** | 0.332 | **0.035** |
| Inc (P) | 0.198 | 0.045 | <0.001 | 0.051 | 0.147 | 0.474 | | 0.253 | 0.251 |
| Inc (A) | <0.001 | 0.073 | <0.001 | <0.001 | 0.084 | 0.051 | 0.075 | | 0.611 |
| Molar | 0.156 | 0.271 | 0.063 | 0.221 | 0.014 | 0.308 | 0.088 | <0.001 | |

**Table 6:** Correlation structure (lower triangle) of *Pseudomys occidentalis* (subsurface); Mantel test $p$-values (upper triangle).

| | Cond | Cond (D) | Cond (P) | Ptery (I) | Mass (S) | Ramus | Inc (P) | Inc (A) | Molar |
|---|---|---|---|---|---|---|---|---|---|
| Cond | | **<0.001** | 0.588 | 0.097 | **0.046** | **0.041** | **0.026** | 0.061 | 0.084 |
| Cond (D) | 0.269 | | 0.246 | 0.056 | 0.231 | **0.013** | 0.492 | 0.955 | **0.035** |
| Cond (P) | <0.001 | 0.034 | | **0.003** | **0.011** | 0.384 | **0.004** | 0.750 | **<0.001** |
| Ptery (I) | 0.077 | 0.111 | 0.288 | | **<0.001** | 0.129 | **<0.001** | 0.625 | **0.004** |
| Mass (S) | 0.103 | 0.044 | 0.222 | 0.315 | | **0.005** | **<0.001** | 0.211 | 0.059 |
| Ramus | 0.089 | 0.147 | 0.011 | 0.077 | 0.188 | | **<0.001** | 0.281 | 0.117 |
| Inc (P) | 0.117 | <0.001 | 0.224 | 0.269 | 0.380 | 0.343 | | 0.099 | **0.042** |
| Inc (A) | 0.094 | <0.001 | <0.001 | <0.001 | 0.051 | 0.027 | 0.094 | | 0.319 |
| Molar | 0.073 | 0.126 | 0.284 | 0.215 | 0.110 | 0.075 | 0.124 | 0.026 | |

**Table 7:** Correlation structure (lower triangle) of *Leporillus apicalis*; Mantel test $p$-values (upper triangle).

| | Cond | Cond (D) | Cond (P) | Ptery (I) | Mass (S) | Ramus | Inc (P) | Inc (A) | Molar |
|---|---|---|---|---|---|---|---|---|---|
| Cond | | **0.008** | 0.177 | 0.481 | 0.368 | 0.307 | 0.246 | 0.301 | 0.650 |
| Cond (D) | 0.248 | | 0.256 | 0.376 | **0.024** | 0.606 | 0.109 | 0.616 | 0.625 |
| Cond (P) | 0.062 | 0.041 | | **0.003** | 0.086 | **0.004** | **0.003** | 0.052 | 0.084 |
| Ptery (I) | <0.001 | 0.010 | 0.265 | | **0.019** | **0.026** | 0.096 | 0.854 | 0.541 |
| Mass (S) | 0.017 | 0.158 | 0.081 | 0.173 | | 0.069 | **0.010** | 0.816 | 0.116 |
| Ramus | 0.027 | <0.001 | 0.288 | 0.218 | 0.123 | | **<0.001** | 0.412 | 0.238 |
| Inc (P) | 0.051 | 0.109 | 0.253 | 0.119 | 0.193 | 0.545 | | 0.073 | **0.010** |
| Inc (A) | 0.028 | <0.001 | 0.089 | <0.001 | <0.001 | 0.006 | 0.099 | | 0.073 |
| Molar | <0.001 | <0.001 | 0.093 | <0.001 | 0.075 | 0.043 | 0.225 | 0.078 | |

**Table 8:** Correlation structure (lower triangle) of *Pseudomys australis*; Mantel test $p$-values (upper triangle).

| | Cond | Cond (D) | Cond (P) | Ptery (I) | Mass (S) | Ramus | Inc (P) | Inc (A) | Molar |
|---|---|---|---|---|---|---|---|---|---|
| Cond | | 0.332 | 0.906 | 0.899 | 0.747 | 0.308 | 0.734 | 0.848 | 0.115 |
| Cond (D) | 0.025 | | **0.005** | 0.060 | 0.349 | 0.172 | **0.038** | 0.705 | 0.410 |
| Cond (P) | <0.001 | 0.194 | | **<0.001** | **<0.001** | 0.311 | **0.030** | 0.902 | 0.499 |
| Ptery (I) | <0.001 | 0.134 | 0.269 | | **0.002** | 0.309 | 0.094 | 0.504 | 0.925 |
| Mass (S) | <0.001 | 0.022 | 0.151 | 0.242 | | 0.110 | **0.008** | 0.131 | 0.774 |
| Ramus | 0.023 | 0.068 | 0.025 | 0.034 | 0.088 | | **0.004** | 0.773 | 0.141 |
| Inc (P) | <0.001 | 0.177 | 0.152 | 0.126 | 0.215 | 0.312 | | 0.059 | 0.429 |
| Inc (A) | <0.001 | <0.001 | <0.001 | <0.001 | 0.082 | <0.001 | 0.142 | | 0.414 |
| Molar | 0.077 | 0.011 | <0.001 | <0.001 | <0.001 | 0.074 | 0.004 | 0.011 | |

**Table 9:** Correlation structure (lower triangle) of *Pseudomys bolami*; Mantel test $p$-values (upper triangle).

| | Cond | Cond (D) | Cond (P) | Ptery (I) | Mass (S) | Ramus | Inc (P) | Inc (A) | Molar |
|---|---|---|---|---|---|---|---|---|---|
| Cond | | 0.501 | **0.041** | **0.028** | 0.097 | 0.999 | 0.133 | 0.730 | 0.810 |
| Cond (D) | <0.001 | | 0.798 | 0.534 | 0.345 | 0.262 | 0.276 | 0.166 | 0.123 |
| Cond (P) | 0.258 | <0.001 | | **<0.001** | 0.258 | 0.649 | **0.049** | 0.983 | 0.565 |
| Ptery (I) | 0.302 | <0.001 | 0.461 | | **0.042** | 0.720 | 0.087 | 0.791 | 0.529 |
| Mass (S) | 0.172 | 0.048 | 0.075 | 0.226 | | 0.190 | **0.010** | 0.381 | 0.726 |
| Ramus | <0.001 | 0.072 | <0.001 | <0.001 | 0.087 | | **<0.001** | **0.037** | 0.060 |
| Inc (P) | 0.130 | 0.060 | 0.189 | 0.166 | 0.276 | 0.539 | | 0.168 | 0.486 |
| Inc (A) | <0.001 | 0.131 | <0.001 | <0.001 | 0.028 | 0.187 | 0.115 | | 0.067 |
| Molar | <0.001 | 0.140 | <0.001 | <0.001 | <0.001 | 0.146 | <0.001 | 0.184 | |

**Table 10:** Correlation structure (lower triangle) of *Pseudomys shortridgei*; Mantel test $p$-values (upper triangle).

| | LC | NM | PA | PA* | PO | PO* | LA | PAu | PB | PS |
|---|---|---|---|---|---|---|---|---|---|---|
| LC | | **0.003** | 0.060 | **0.011** | **0.015** | 0.064 | **0.049** | **0.024** | **0.010** | 0.068 |
| NM | 0.446 | | **0.002** | **0.002** | **0.043** | 0.213 | **0.007** | **0.008** | **0.050** | **0.029** |
| PA | 0.273 | 0.573 | | **<0.001** | **0.022** | 0.171 | **<0.001** | **0.003** | **0.003** | **0.018** |
| PA* | 0.450 | 0.592 | 0.587 | | **0.008** | **0.041** | **0.002** | **0.002** | **0.002** | **0.041** |
| PO | 0.430 | 0.272 | 0.382 | 0.518 | | **0.002** | **0.013** | **0.003** | **0.005** | **0.015** |
| PO* | 0.358 | 0.118 | 0.157 | 0.336 | 0.658 | | **0.042** | 0.054 | 0.232 | 0.088 |
| LA | 0.329 | 0.361 | 0.604 | 0.528 | 0.433 | 0.360 | | **0.006** | **0.003** | **0.005** |
| PAu | 0.391 | 0.444 | 0.492 | 0.701 | 0.712 | 0.352 | 0.512 | | **0.005** | **0.010** |
| PB | 0.458 | 0.275 | 0.537 | 0.529 | 0.467 | 0.171 | 0.553 | 0.539 | | **0.005** |
| PS | 0.247 | 0.373 | 0.369 | 0.325 | 0.383 | 0.213 | 0.445 | 0.427 | 0.438 | |

**Table 11:** Pairwise correlations between samples (lower triangle) and Mantel test $p$-values (upper triangle). Abbreviations: LC = *Leporillus conditor*, NM = *Notomys mitchelli*, PA = *Pseudomys albocinereus*, PA* = *P. albocinereus* (subsurface), PO = *P. occidentalis*, PO* = *P. occidentalis* (subsurface), LA = *L. apicalis*, PAu = *P. australis*, PB = *P. bolami*, PS = *P. shortridgei*.

| | LC | NM | PA | PA* | PO | PO* | LA | PAu | PB | PS |
|---|---|---|---|---|---|---|---|---|---|---|
| LC | 0.508 | 0.761 | **0.010** | 0.530 | 0.228 | 0.658 | **0.019** | 0.114 | 0.480 | 0.162 |
| NM | 0.851 | 0.581 | 0.443 | 0.879 | **0.036** | 0.144 | **0.038** | 0.197 | 0.101 | 0.422 |
| PA | 0.527 | 0.945 | 0.762 | 0.893 | 0.130 | 0.201 | 0.558 | 0.336 | 0.713 | 0.442 |
| PA* | 0.864 | 0.966 | 0.496 | 0.648 | 0.474 | 0.595 | 0.304 | 0.927 | 0.678 | 0.307 |
| PO | 0.860 | 0.292 | **0.045** | 0.737 | 0.720 | 0.993 | 0.089 | 0.948 | 0.520 | 0.468 |
| PO* | 0.688 | 0.076 | **<0.001** | 0.282 | 0.890 | 0.535 | **0.026** | 0.070 | **0.023** | 0.098 |
| LA | 0.649 | 0.547 | 0.554 | 0.788 | 0.246 | 0.650 | 0.764 | 0.365 | 0.776 | 0.558 |
| PAu | 0.771 | 0.774 | 0.204 | 0.977 | 0.947 | 0.654 | 0.261 | 0.734 | 0.735 | 0.529 |
| PB | 0.878 | 0.314 | 0.349 | 0.767 | 0.316 | 0.247 | 0.351 | 0.482 | 0.674 | 0.572 |
| PS | 0.477 | 0.580 | **0.028** | 0.252 | 0.123 | 0.316 | 0.129 | 0.163 | 0.417 | 0.635 |

**Table 12:** Significances of testing whether two correlation structures are no more different than by chance (inversion of the null hypothesis of the Mantel test); lower and upper triangles correspond to $p$-values derived from the correlation structure permuted by rows and columns, respectively. Diagonals are matrix repeatabilities of sample correlation matrices. See Table 11 for abbreviations.

| | LC | NM | PA | PA* | PO | PO* | LA | PAu | PB |
|---|---|---|---|---|---|---|---|---|---|
| NM | 1.757 | | | | | | | | |
| PA | 2.123 | 1.228 | | | | | | | |
| PA* | 1.858 | 1.369 | 0.782 | | | | | | |
| PO | 2.439 | 1.467 | 1.008 | 1.522 | | | | | |
| PO* | 1.665 | 1.218 | 1.044 | 0.903 | 1.402 | | | | |
| LA | 1.995 | 1.736 | 1.703 | 1.739 | 1.761 | 1.903 | | | |
| PAu | 2.348 | 1.247 | 0.823 | 1.233 | 0.989 | 1.266 | 1.838 | | |
| PB | 1.866 | 1.063 | 1.232 | 1.503 | 1.170 | 1.117 | 1.809 | 0.942 | |
| PS | 2.729 | 2.008 | 1.741 | 2.291 | 1.538 | 2.251 | 1.858 | 1.594 | 1.745 |

**Table 13:** Pairwise distances between samples in the PCoA space resulting from relative eigenanalysis. See Table 11 for abbreviations.

| | LC | NM | PA | PA* | PO | PO* | LA | PAu | PB |
|---|---|---|---|---|---|---|---|---|---|
| NM | 9.800 | | | | | | | | |
| PA | 9.807 | 7.007 | | | | | | | |
| PA* | 9.807 | 7.007 | 0.016 | | | | | | |
| PO | 9.807 | 7.007 | 6.414 | 6.414 | | | | | |
| PO* | 9.807 | 7.007 | 6.414 | 6.414 | 0.016 | | | | |
| LA | 4.400 | 9.800 | 9.807 | 9.807 | 9.807 | 9.807 | | | |
| PAu | 9.800 | 7.000 | 5.607 | 5.607 | 6.407 | 6.407 | 9.800 | | |
| PB | 9.800 | 7.000 | 5.607 | 5.607 | 6.407 | 6.407 | 9.800 | 4.500 | |
| PS | 9.800 | 7.000 | 5.607 | 5.607 | 6.407 | 6.407 | 9.800 | 5.200 | 5.200 |

**Table 14:** Table of phylogenetic distances in millions of years, derived from phylogeny by Roycroft *et al.* (2021). See Table 11 for abbreviations.

| | Cond | Cond (D) | Cond (P) | Ptery (I) | Mass (S) | Ramus | Inc (P) | Inc (A) | Molar |
|---|---|---|---|---|---|---|---|---|---|
| Cond | | 1.000 | 0.093 | 0.469 | **0.023** | 1.000 | 1.000 | 0.315 | 1.000 |
| Cond (D) | <0.001 | | 1.000 | 0.065 | 1.000 | 1.000 | 1.000 | 1.000 | 1.000 |
| Cond (P) | 0.961 | <0.001 | | 1.000 | 0.108 | 1.000 | 1.000 | 0.110 | 1.000 |
| Ptery (I) | 0.587 | 0.986 | <0.001 | | 1.000 | 1.000 | 1.000 | 1.000 | 1.000 |
| Mass (S) | 0.911 | <0.001 | 0.950 | <0.001 | | 1.000 | 1.000 | 1.000 | 1.000 |
| Ramus | <0.001 | <0.001 | <0.001 | <0.001 | <0.001 | | 1.000 | **0.001** | 1.000 |
| Inc (P) | <0.001 | <0.001 | <0.001 | <0.001 | <0.001 | <0.001 | | **0.046** | 1.000 |
| Inc (A) | 0.659 | <0.001 | 0.664 | <0.001 | <0.001 | 0.999 | 0.994 | | 0.346 |
| Molar | <0.001 | <0.001 | <0.001 | <0.001 | <0.001 | <0.001 | <0.001 | 0.831 | |

**Table 15:** Pagel's $\lambda$ for partial correlations between partitions (lower triangle); $p$-values based on log-ratio tests (upper triangle), significant values bolded.

| | Cond | Cond (D) | Cond (P) | Ptery (I) | Mass (S) | Ramus | Inc (P) | Inc (A) | Molar |
|---|---|---|---|---|---|---|---|---|---|
| Cond | | 0.116 | **0.039** | 0.237 | **0.034** | 0.938 | 0.811 | 0.289 | 0.955 |
| Cond (D) | 0.061 | | 0.279 | **0.014** | 0.956 | 0.523 | 0.224 | 0.822 | 0.906 |
| Cond (P) | 0.152 | 0.027 | | 0.384 | 0.070 | 0.452 | 0.465 | 0.222 | 0.736 |
| Ptery (I) | 0.028 | 0.275 | 0.023 | | 0.070 | 0.837 | 0.781 | 1.000 | 0.829 |
| Mass (S) | 0.119 | 0.003 | 0.128 | 0.096 | | 0.382 | 0.689 | 0.538 | 0.778 |
| Ramus | 0.004 | 0.012 | 0.010 | 0.005 | 0.014 | | 0.422 | **0.006** | 0.932 |
| Inc (P) | 0.006 | 0.027 | 0.017 | 0.005 | 0.007 | 0.015 | | **0.017** | 0.532 |
| Inc (A) | 0.021 | 0.005 | 0.027 | 0.002 | 0.011 | 0.934 | 0.395 | | 0.098 |
| Molar | 0.003 | 0.004 | 0.007 | 0.006 | 0.006 | 0.004 | 0.011 | 0.057 | |

**Table 16:** Blomberg *et al.*'s $K$ for partial correlations between partitions (lower triangle); $p$-values based on 1000 permutations (upper triangle), significant values bolded.

| | | Ward's | UPGMA |
|---|---|---|---|
| **Correlations** | Cophenetic correlation | 0.480 | 0.704 |
| | Agglomerative coefficient | **0.498** | 0.334 |
| **PCoA Distances** | Cophenetic correlation | 0.839 | **0.879** |
| | Agglomerative coefficient | **0.472** | 0.400 |

**Table 17:** Clustering statistics for Ward's and UPGMA clustering of the integration disparity matrix (top rows) and distances in the PCoA space resulting from relative eigenanalysis (bottom rows). Significant values bolded. UPGMA clustering shown in Figure 19.

### 7 Supplemental Figures

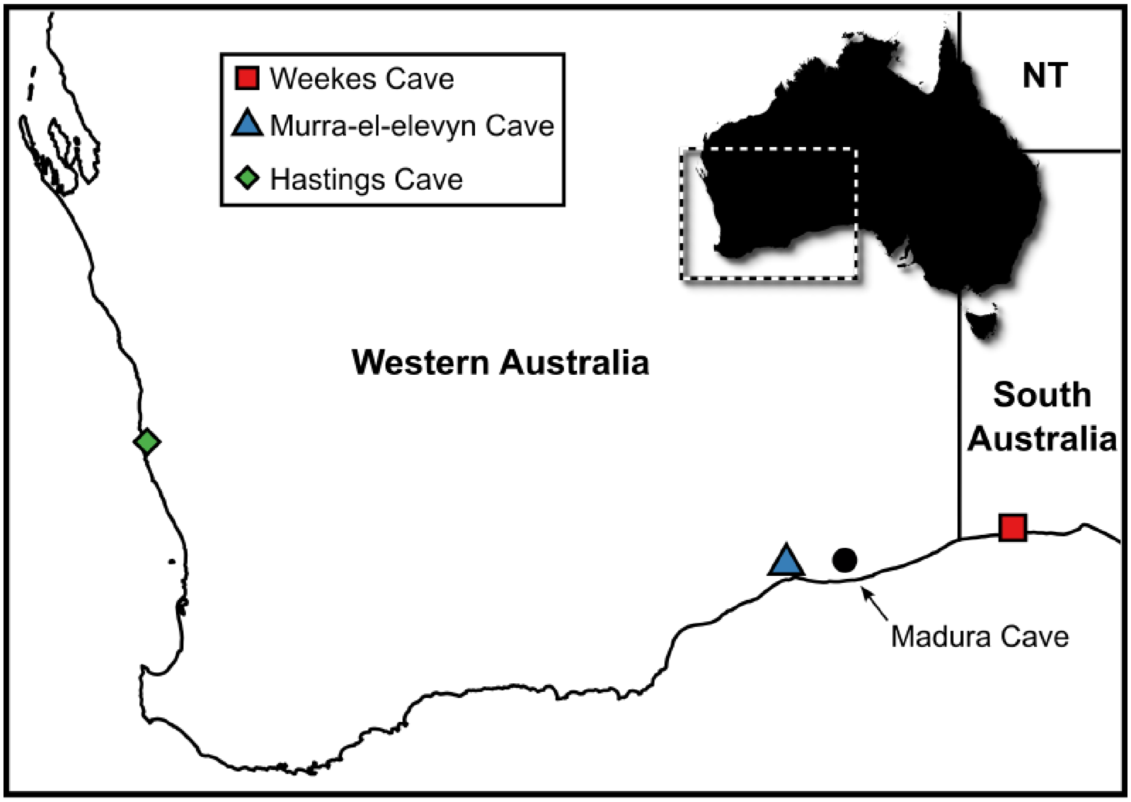

**Figure 1:** Locality map including the well-studied Madura Cave.

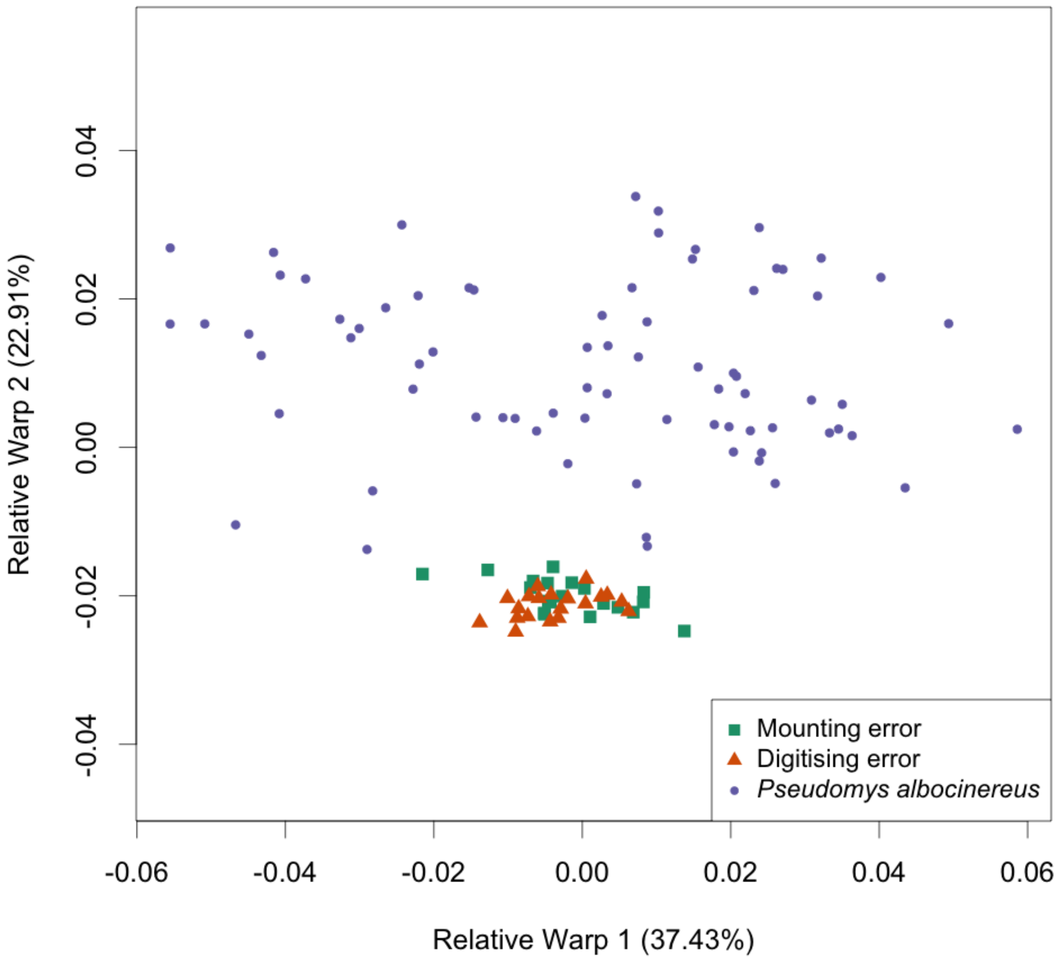

**Figure 2:** Morphospace of surface sample of *Pseudomys albocinereus* and an assessment of mounting and digitising error.

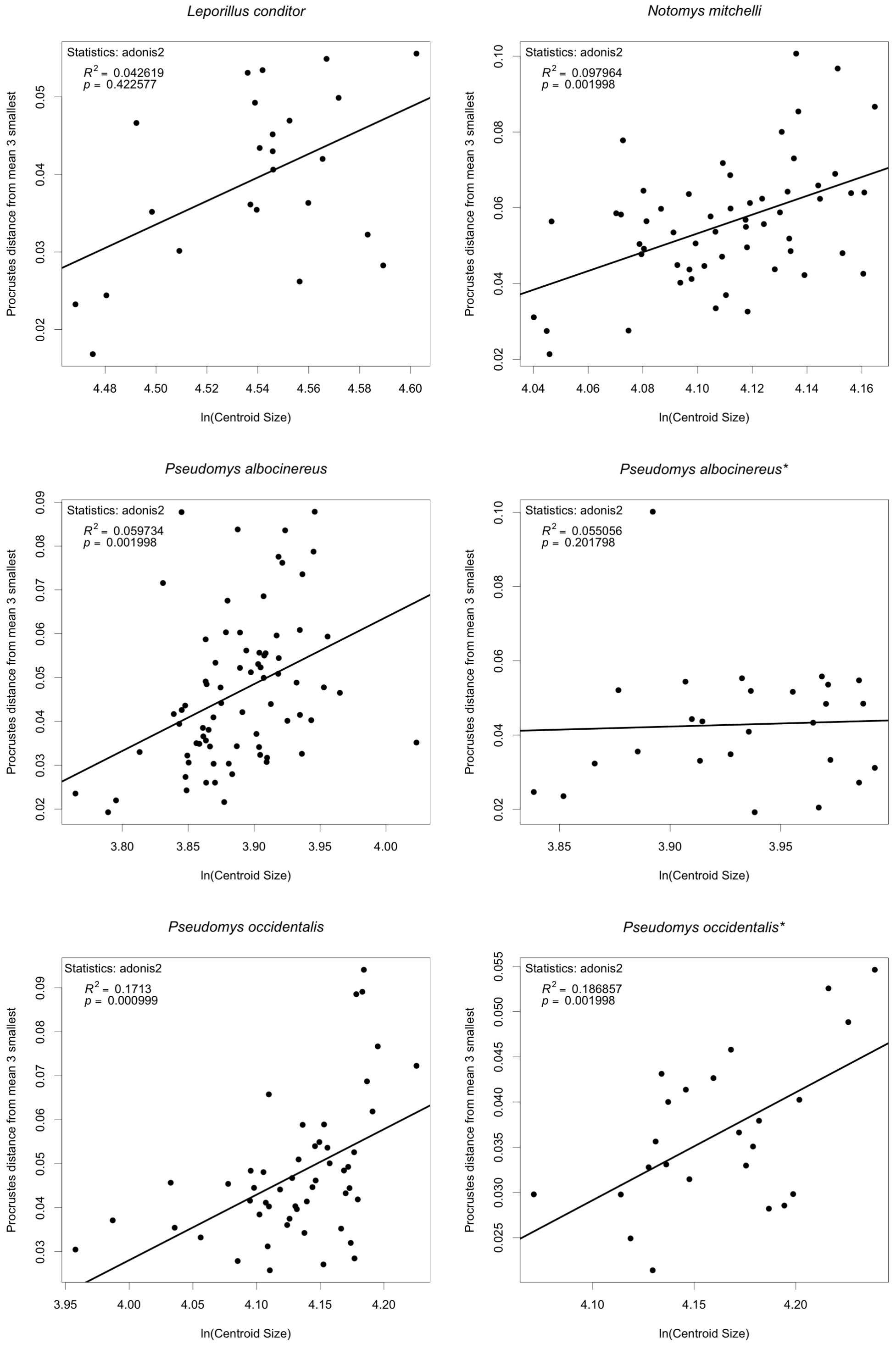

**Figure 3:** Allometry; the natural logarithm of centroid size versus Procrustes distance from the mean of the three smallest specimens for each sample (continued in Figure 4). Some relationships are significant but very weak. Starred taxa are subsurface samples.

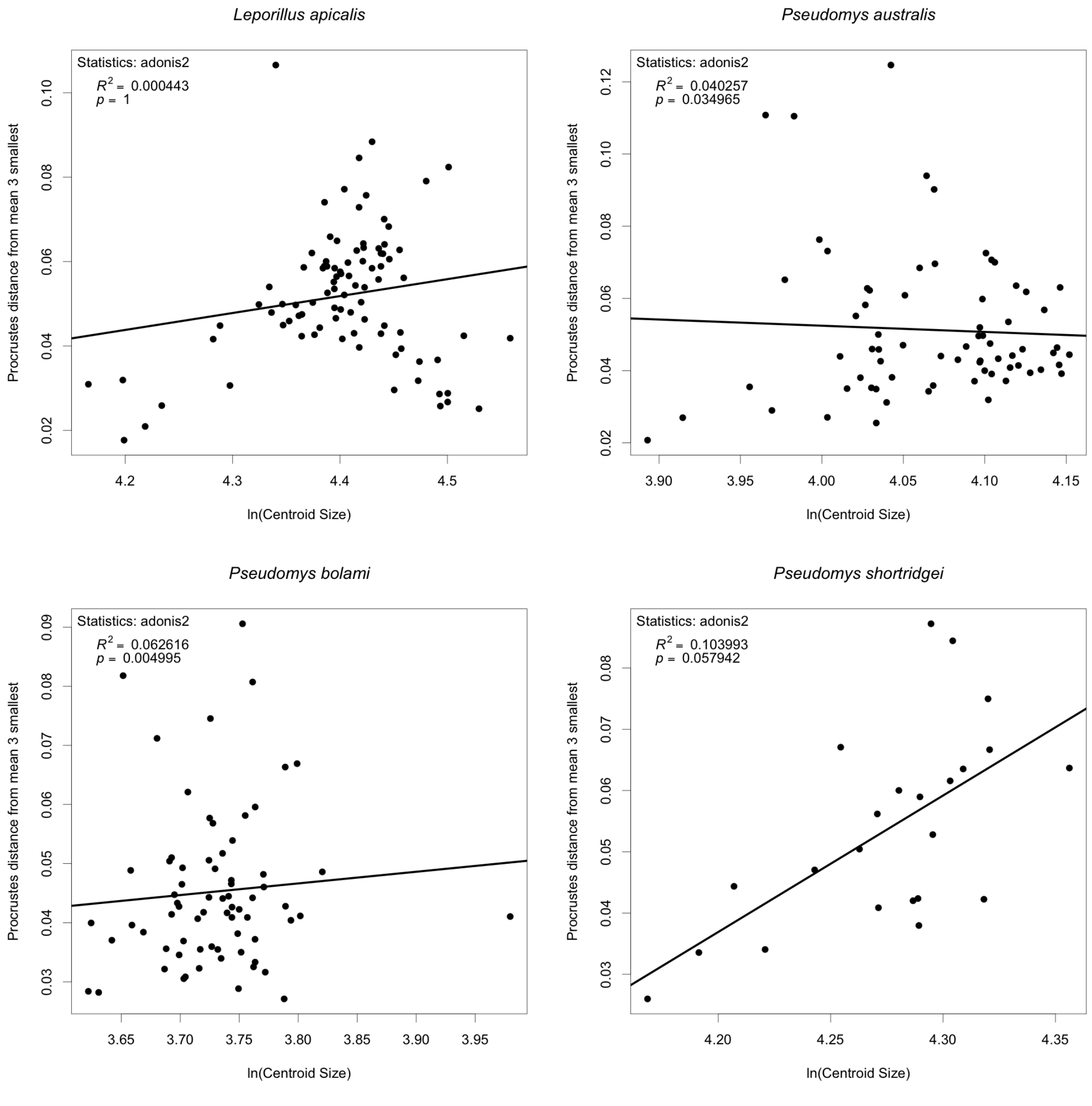

**Figure 4:** Continued from Figure 3.

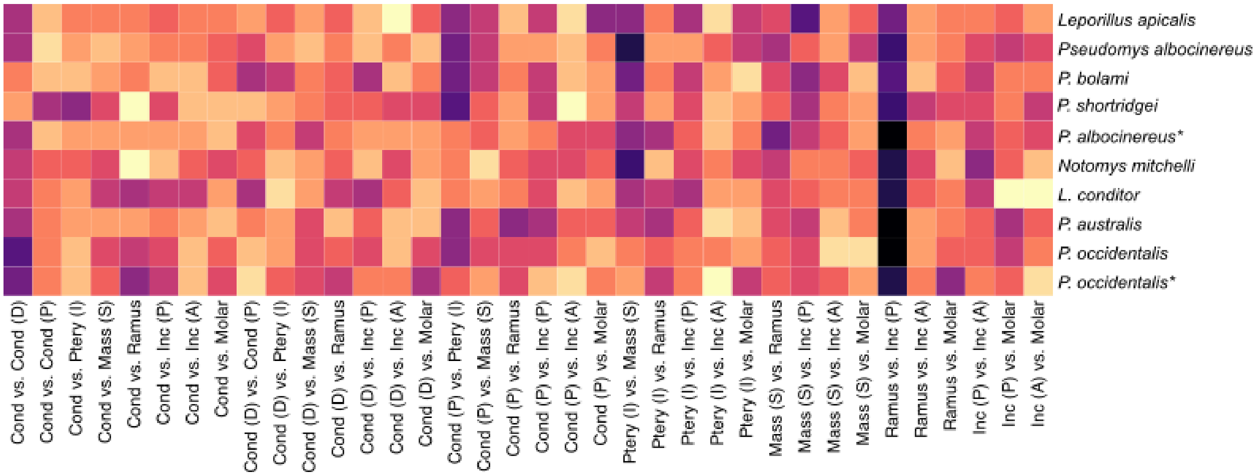

**Figure 5:** Heatmap comparison of correlation structures of all samples studied. Colour scale is relative between samples; darker colours are stronger correlations. Starred taxa are subsurface samples. Sample rows organised according to Ward's clustering of the integration disparity matrix. See Table 1 through Table 10 and Figure 6 through Figure 15 for a complete report of each correlation structure. Starred taxa are subsurface samples.

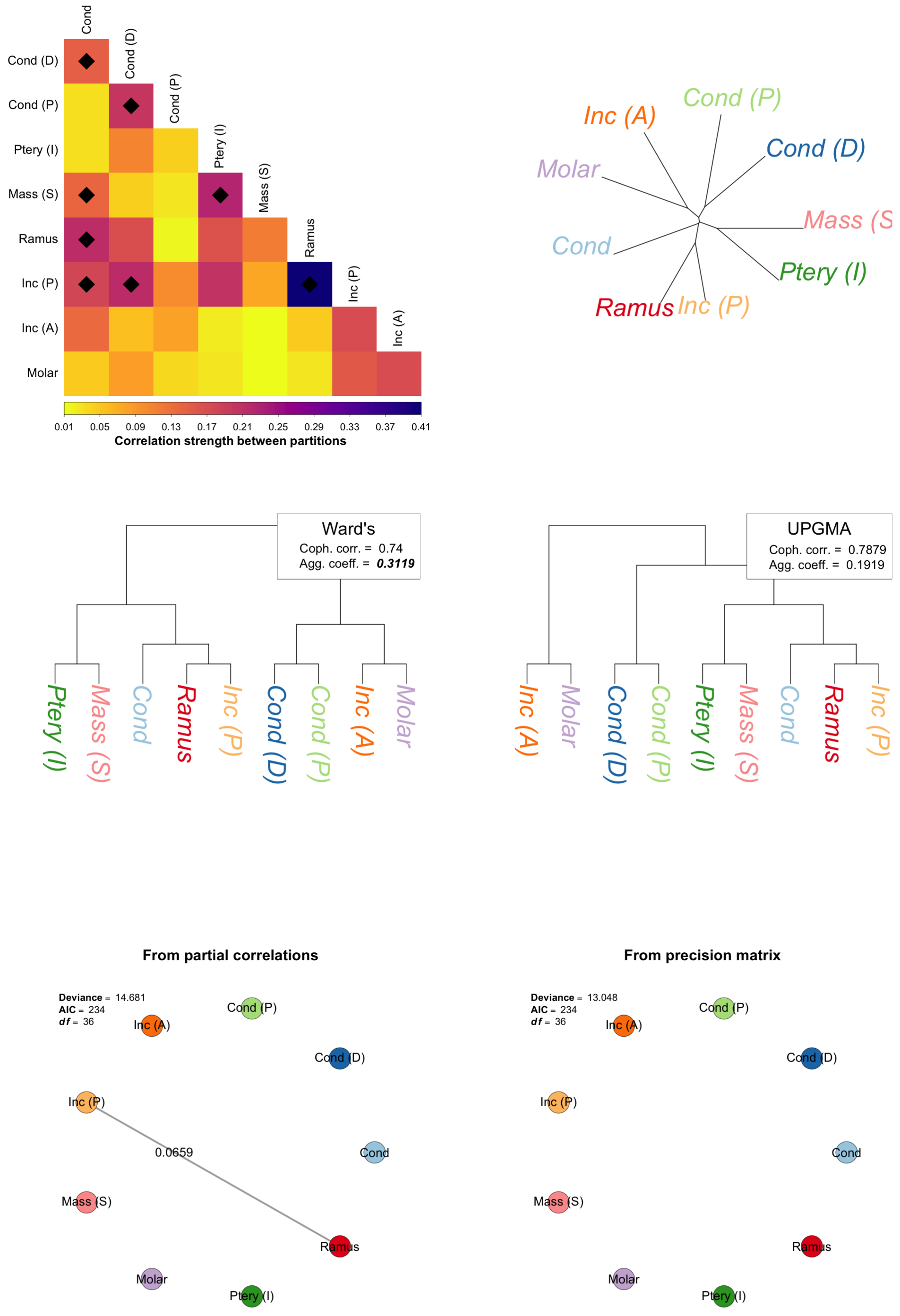

**Figure 6:** Correlation structure of *Leporillus conditor* and associated exploratory analyses. **Top row:** left, correlation structure as heatmap with significant correlations starred (see tables for exact values); right, reticulate network analysis with reticulations (if present) indicated in blue dashed lines. **Middle row:** left, Ward's clustering of correlation structure with associated cophenetic correlation and agglomerative coefficient (bolded for significance); right, UPGMA clustering with statistics denoted as at left. **Bottom row:** left, graphical modelling based on partial correlations between partitions with associated model statistics and weak edges denoted by dashed lines; right, graphical modelling based off of the precision matrix (full set of correlations) with statistics and edges denoted as at left.

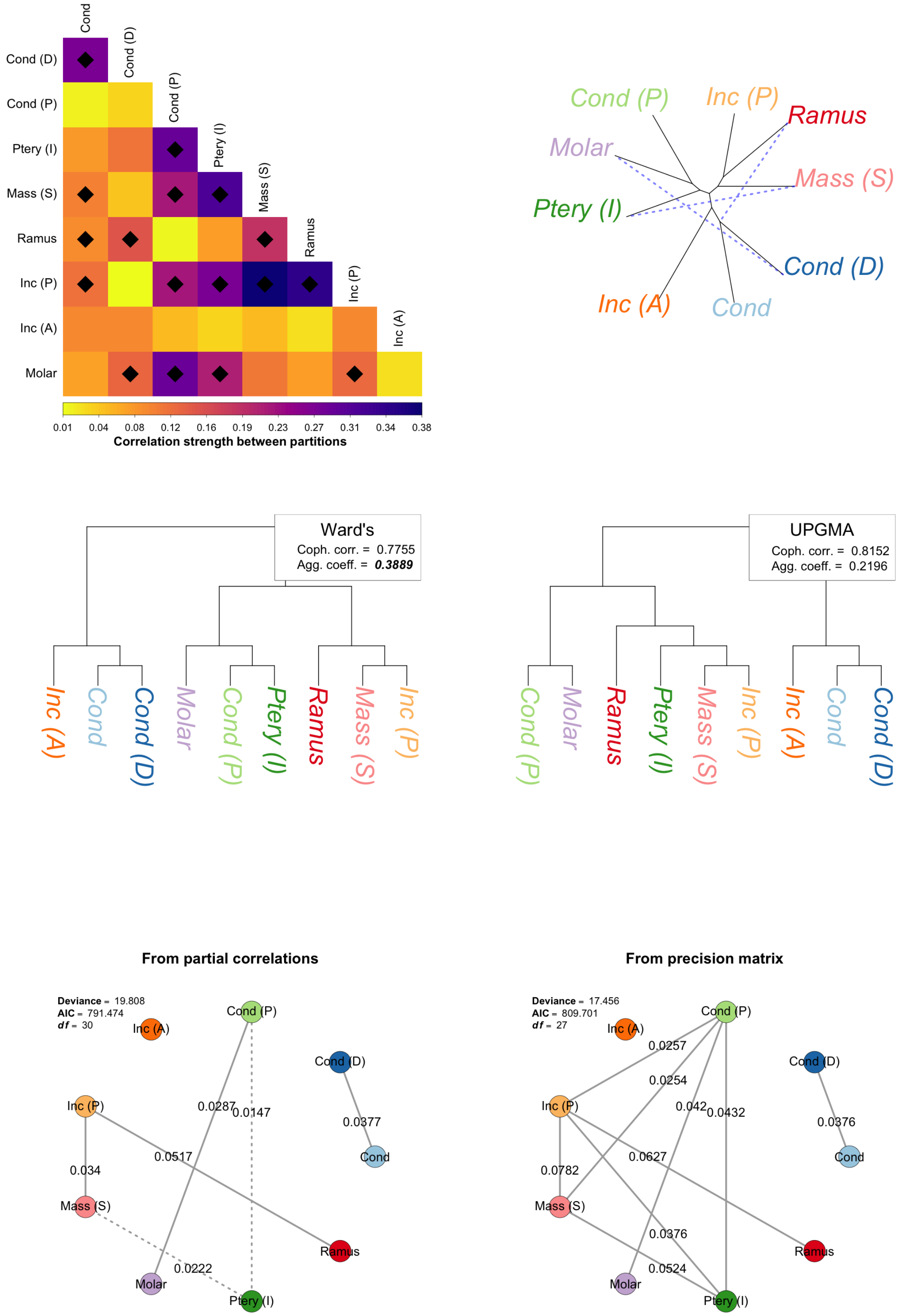

**Figure 7:** Correlation structure of *Leporillus apicalis* and associated exploratory analyses. See caption to Figure 6 for explanation.

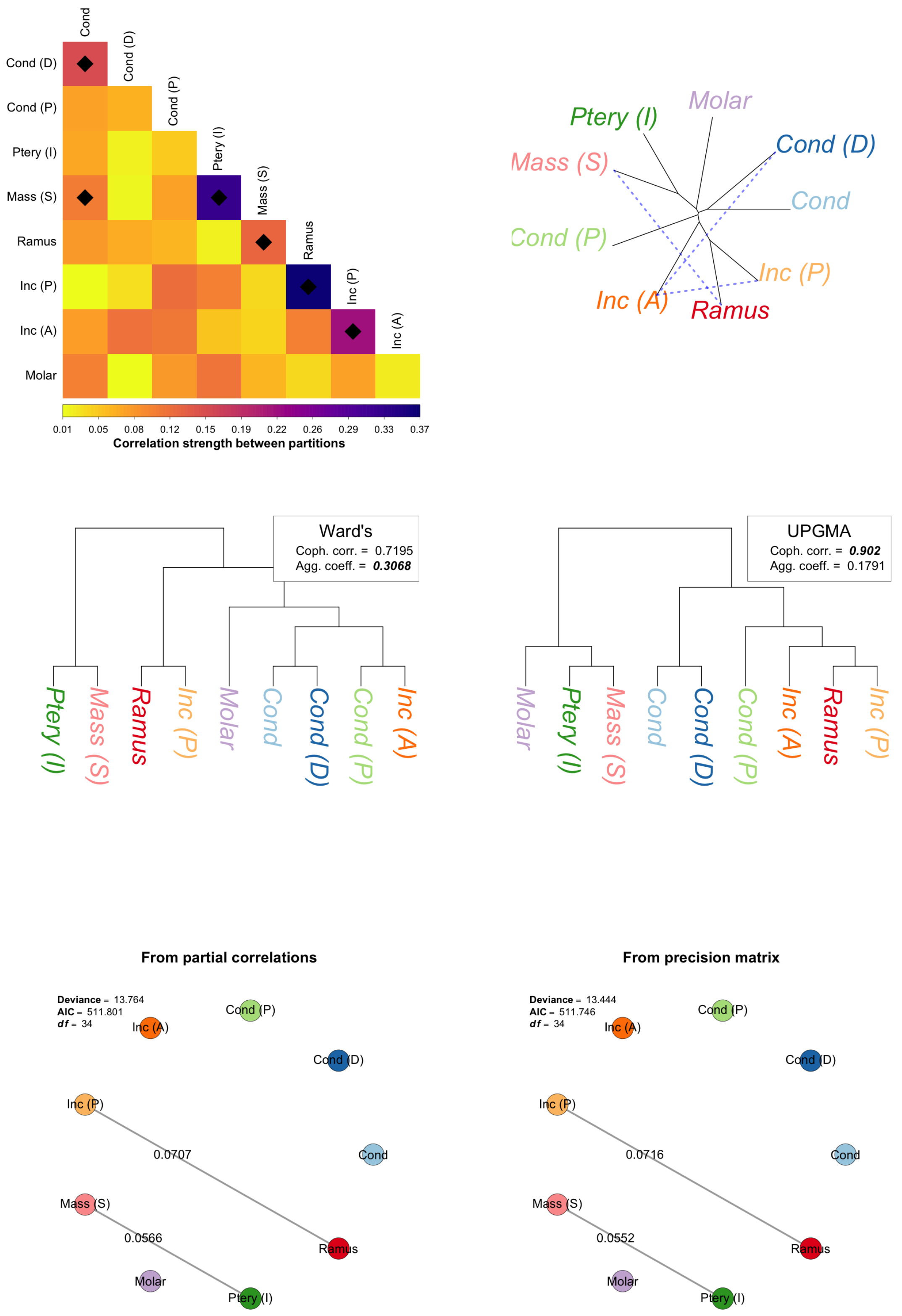

**Figure 8:** Correlation structure of *Notomys mitchelli* and associated exploratory analyses. See caption to Figure 6 for explanation.

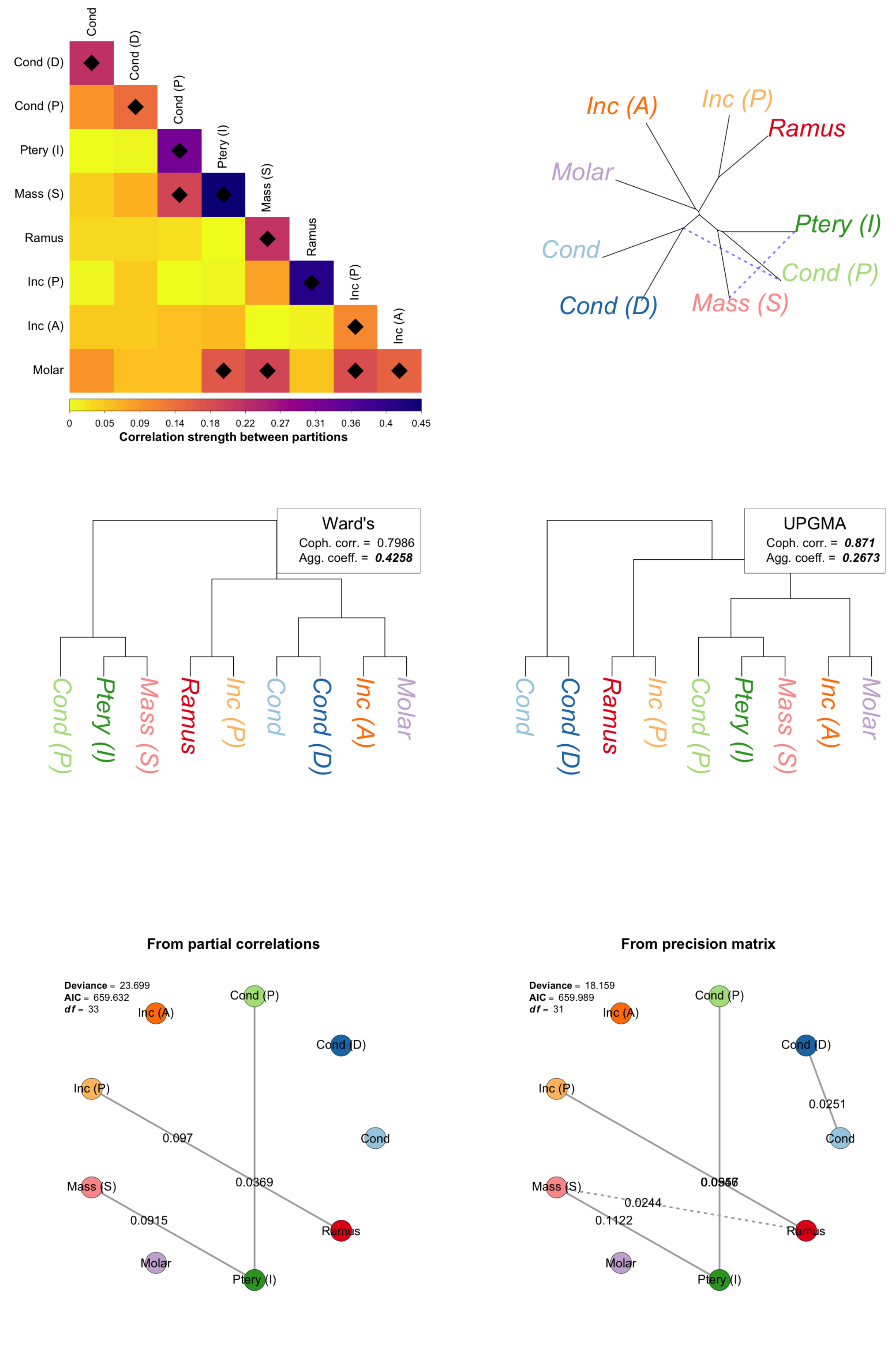

**Figure 9:** Correlation structure of *Pseudomys albocinereus* (surface sample) and associated exploratory analyses. See caption to Figure 6 for explanation.

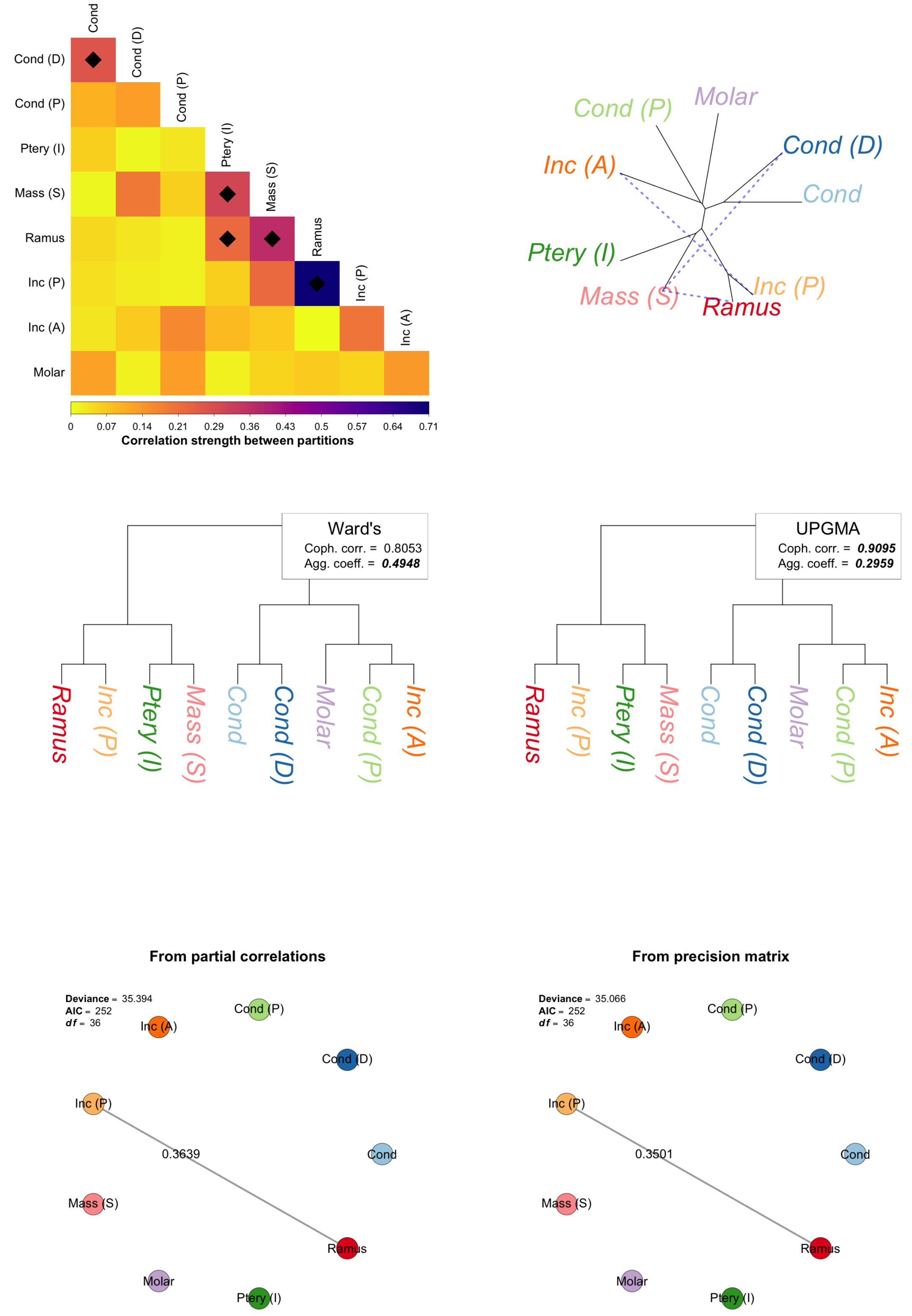

**Figure 10:** Correlation structure of *Pseudomys albocinereus* (subsurface sample) and associated exploratory analyses. See caption to Figure 6 for explanation.

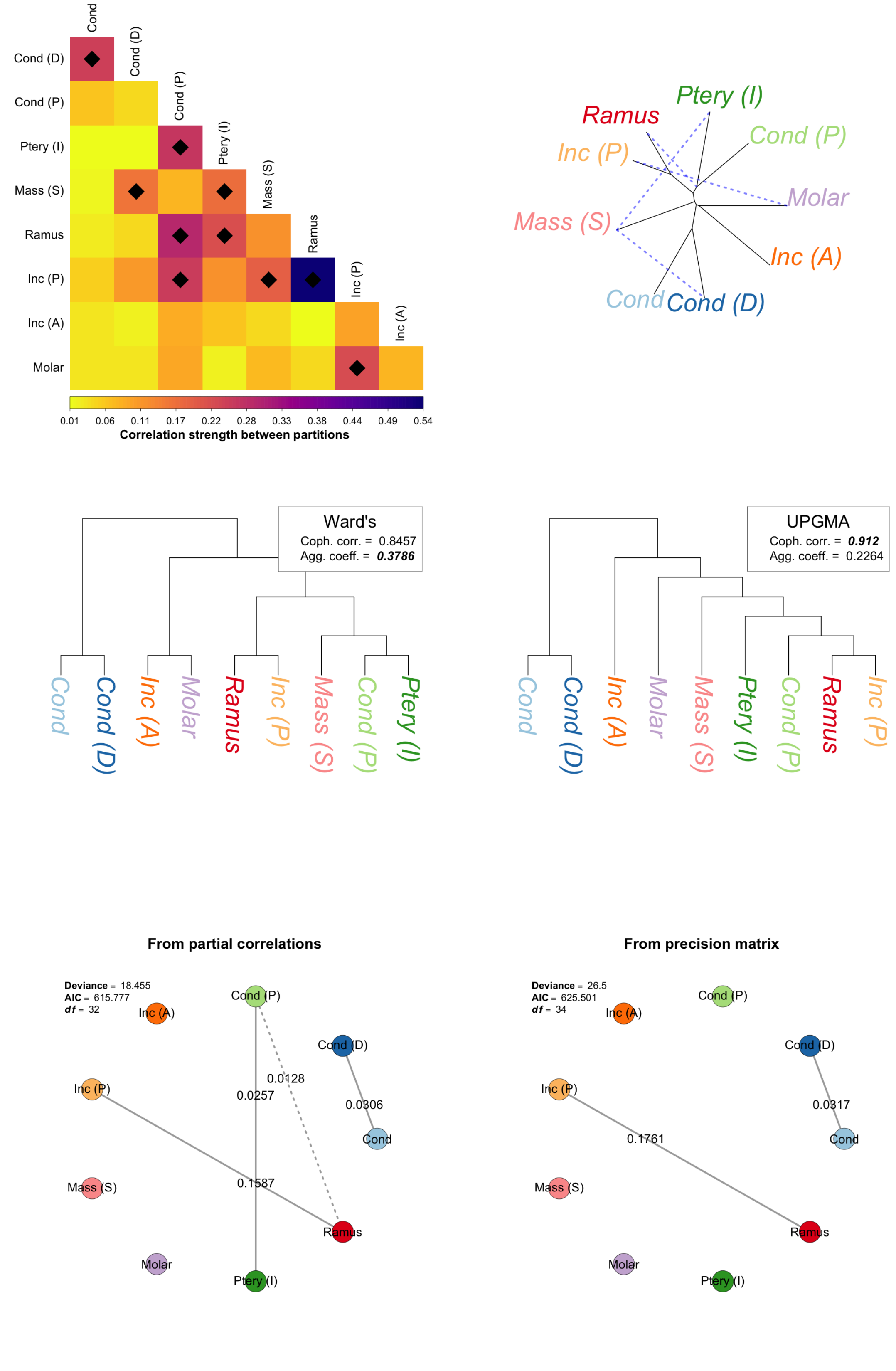

**Figure 11:** Correlation structure of *Pseudomys australis* and associated exploratory analyses. See caption to Figure 6 for explanation.

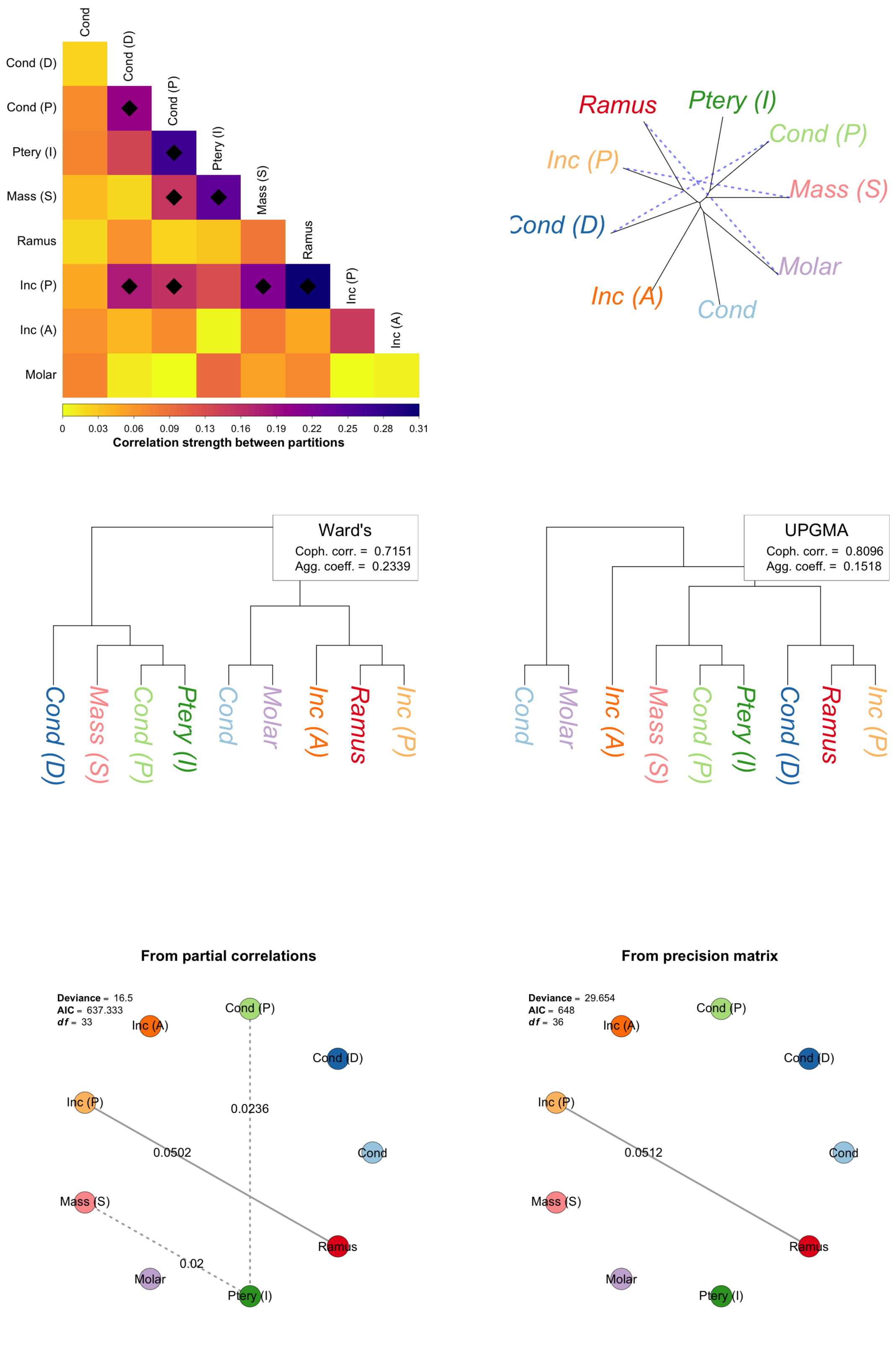

**Figure 12:** Correlation structure of *Pseudomys bolami* and associated exploratory analyses. See caption to Figure 6 for explanation.

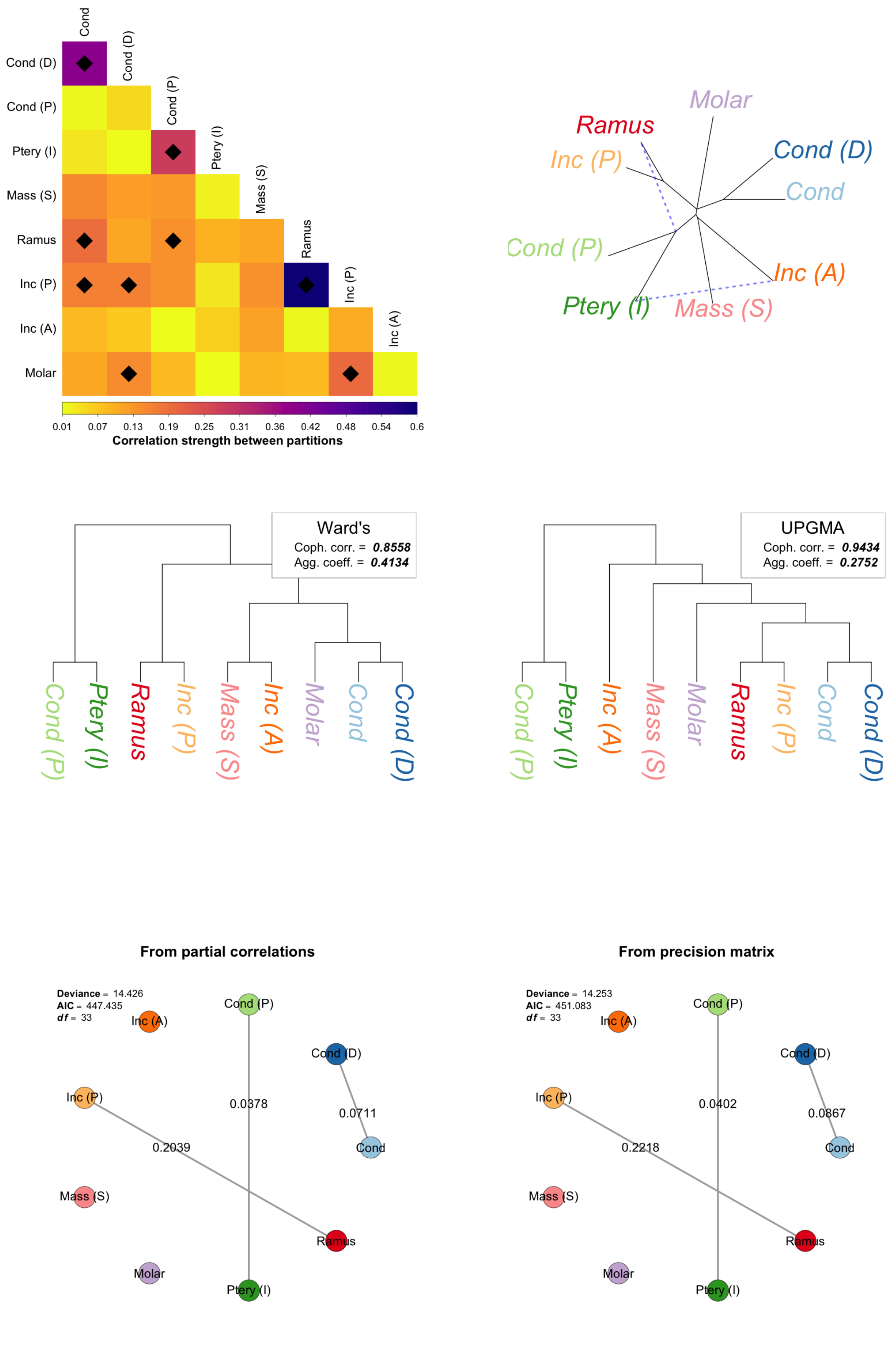

**Figure 13:** Correlation structure of *Pseudomys occidentalis* (surface sample) and associated exploratory analyses. See caption to Figure 6 for explanation.

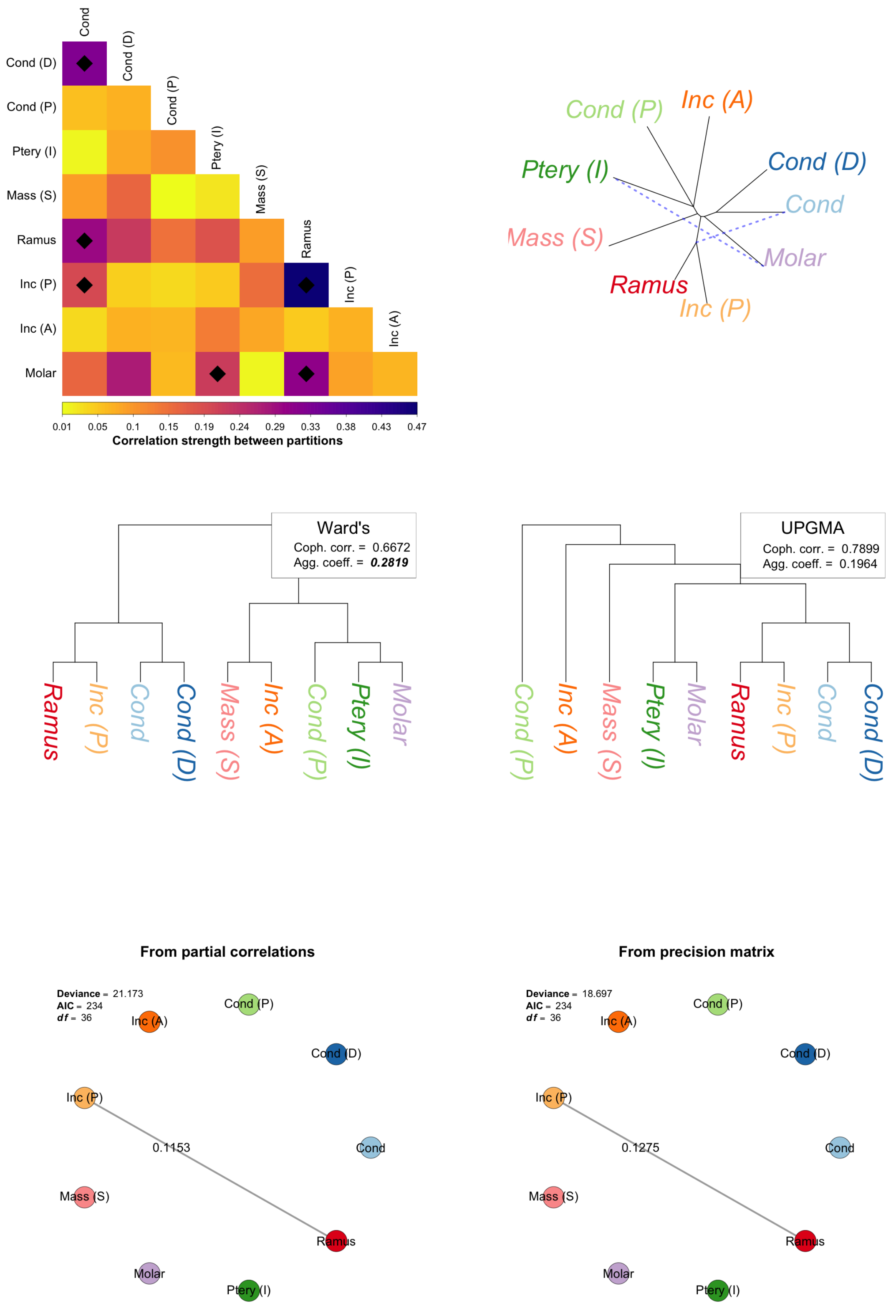

**Figure 14:** Correlation structure of *Pseudomys occidentalis* (subsurface sample) and associated exploratory analyses. See caption to Figure 6 for explanation.

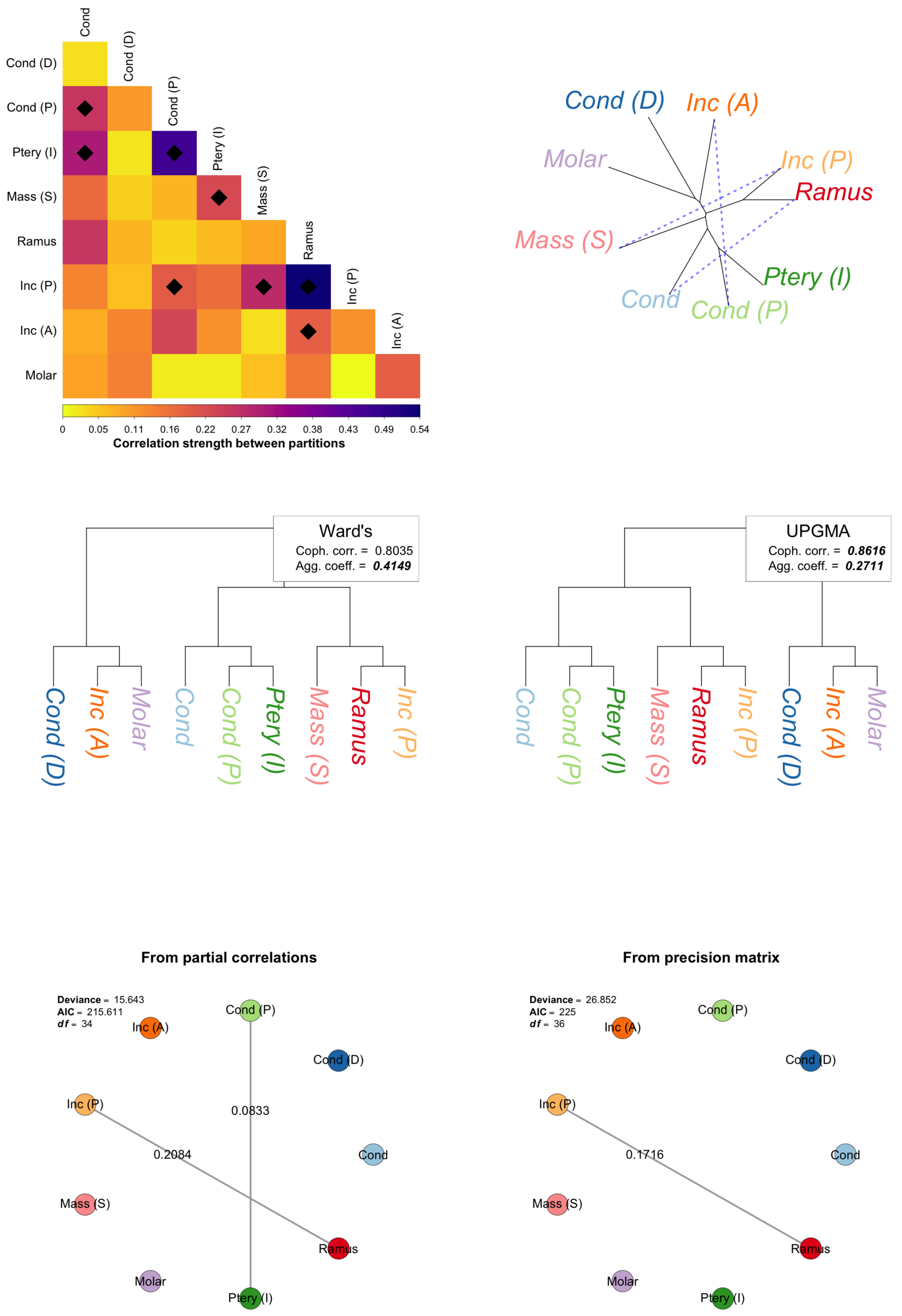

**Figure 15:** Correlation structure of *Pseudomys shortridgei* and associated exploratory analyses. See caption to Figure 6 for explanation.

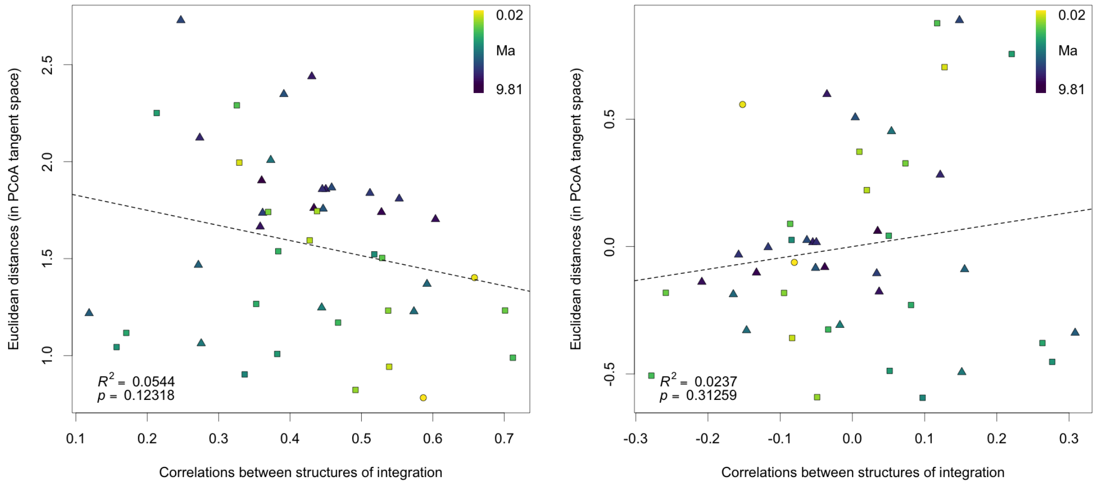

**Figure 16:** Pairwise comparisons between structures of integration compared; partition-based between sample correlations versus distances in the PCoA space derived from relative eigenanalysis. Coloured by degree of phylogenetic separation. Point shapes: circles = conspecifics, squares = congenerics, triangles = confamilials. Left: raw comparison. Right: comparison of residuals after correcting for phylogenetic signal.

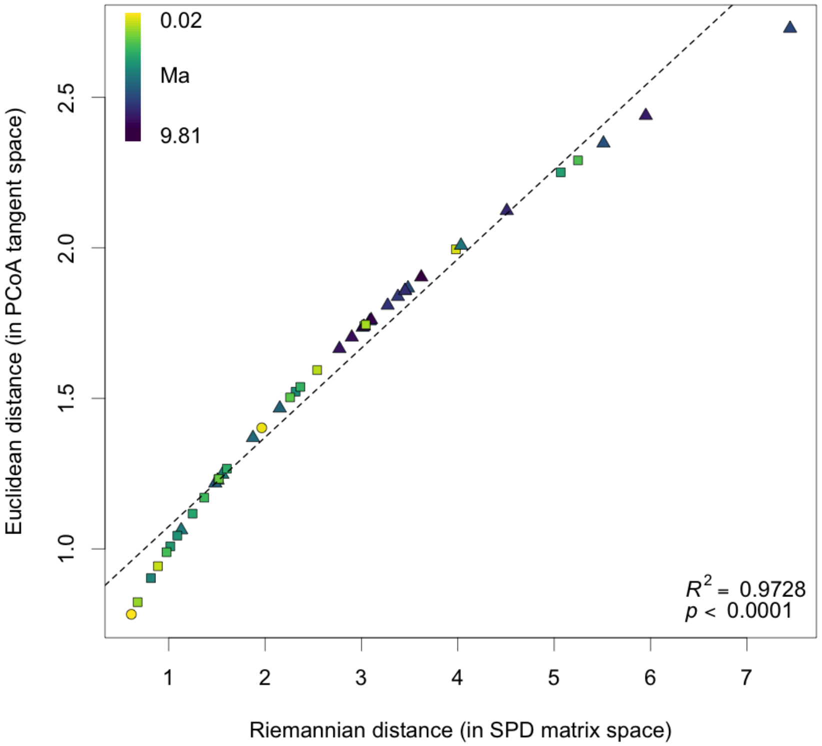

**Figure 17:** Minimal distortion by projection of Riemannian distances into Euclidean space in the course of relative eigenanalysis. Colours and points as in Figure 16.

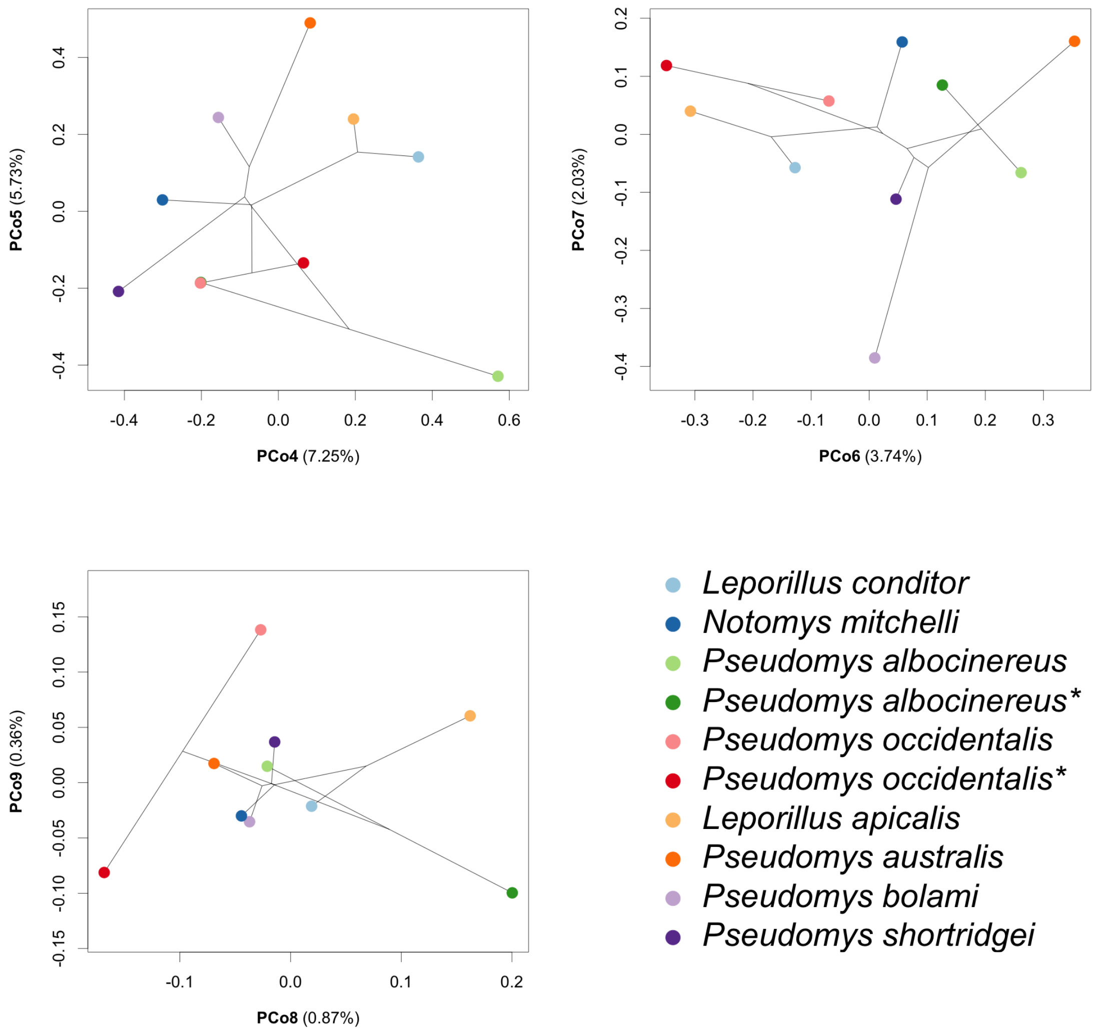

**Figure 18:** Relative eigenanalysis results: higher principal coordinate axes. See text for first three axes.

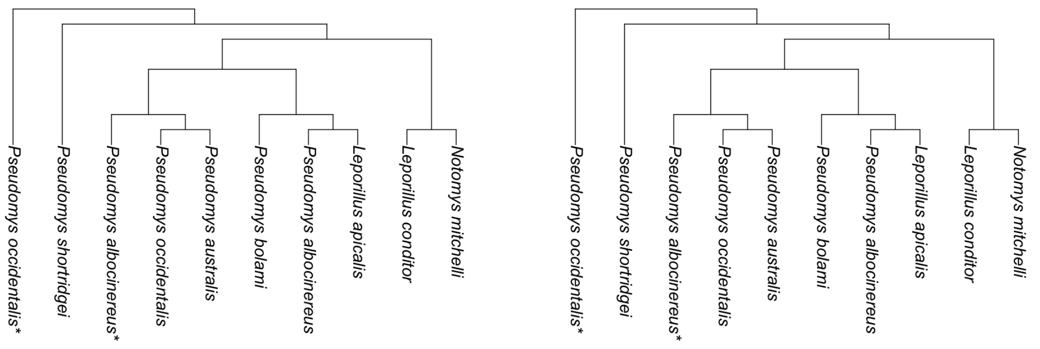

**Figure 19:** UPGMA clustering of integration disparity matrix (left) and distance matrix resulting from relative eigenanalysis (right). See Table 17 for statistics on these clustering methods.
